# Gill regeneration in the mayfly *Cloeon* uncovers new molecular pathways in insect regeneration

**DOI:** 10.1101/2024.04.17.589898

**Authors:** Carlos A. Martin-Blanco, Pablo Navarro, José Esteban-Collado, Florenci Serras, Isabel Almudi, Fernando Casares

## Abstract

The capacity to regenerate lost or damaged organs is widespread among animals, and yet, the species in which regeneration has been experimentally probed using molecular and functional assays is very small. This is also the case for insects, for which we still lack a complete picture of their regeneration mechanisms and the extent of conservation of these mechanisms. Here we contribute to filling this gap by investigating regeneration in the mayfly *Cloeon dipterum.* Mayflies, or Ephemeroptera, appeared early in the evolution of insects. We focus on the abdominal gills of *Cloeon* nymphs, which are critical for osmoregulation and gas exchange. After amputation, gills re-grow faster than they do during normal development. Direct cell count and EdU proliferation assays indicate that growth acceleration involves an uniform increase in cell proliferation throughout the gill, rather than a localized growth zone. Transcriptomic analysis reveals an early enrichment in cell cycle-related genes, in agreement with fast proliferation. Several other gene classes are also enriched in regenerating gills, including protein neddylation and other proteostatic processes. We then showed that protein neddylation, the activin signaling pathway or the mRNA-binding protein Lin28, among other genes and processes, are required for *Drosophila* larval/pupal wing regeneration, and that some of these genes may have a regeneration-specific function in the wing. Globally, our results contribute to elucidating regeneration mechanisms in mayflies and suggest a conservation of regeneration mechanisms across insects, as evidenced by the regenerative role of candidate genes identified in *Cloeon* in the distant *Drosophila*.

## INTRODUCTION

Regeneration allows restoring function, partially or fully, after an organ has been lost or damaged. This capacity is widespread, although scattered, along the animal tree of life, with some groups (or even species within groups) being able to fully regenerate their organs, while others are capable only of partial regeneration, or lack that ability altogether (1–3)

Among animals, crustacea and insects (Pancrustacea) are excellent regenerators, especially of their limbs (4, 5). Specifically within insects, regeneration has been described in species belonging to 38 genera (6). Insects secrete an exoskeleton, so that growth is allowed by the regular shedding, or molting, of the exoskeleton until the animal reaches adulthood. Furthermore, as a general rule, regeneration in insects requires molting. Thus, as adult pterygote insects do not molt, regeneration only occurs during the larval or nymphal period.

Despite the broad distribution of regenerative capacity within insects, the search for molecular and cellular mechanisms involved in insect appendage regeneration has been restricted to a small set of species (reviewed in (4, 7)). This research has identified a number of genetic components and pathways that seem to be shared during limb regeneration in insects such as crickets and cockroaches or flies (8–14), and includes the early involvement of the Jun stress response and the JAK/STAT pathways, the Hippo and Insulin/IGF growth-control pathways, the activation of patterning signaling pathways, such as wg/Wnt, hh/Shh, EGF, dpp/BMP, Notch and Toll, the usage of transcription factors known to be critical in proximo-distal leg patterning, such as *Dll*, *dac* or *hth*, and the requirement of the planar *Ds/ft* polarity pathway (7). However, there are major gaps in our understanding of insect regeneration and the extent to which regenerative mechanisms are conserved across species and organs. Several studies have found that genes and pathways previously known to play a major role during limb development are also involved in limb regeneration, raising the possibility that regeneration recapitulates the developmental program (15). However, the discovery of damage/regeneration-specific regulatory elements suggests that the gene networks responsible for the early phases of wound healing and regeneration differ from development (16–18). Moreover, recent work on the malacostracan species *Parhyale hawaiensis*, (both insects and malacostracans belong to the clade Pancrustacea) shows that, although leg development and regeneration share a similar set of expressed genes, the temporal deployment of these genes differs (19).

The Pond Olive mayfly *Cloeon dipterum* is a freshwater insect which only emerges from the water as a reproductive, flying adult (20). The aquatic juveniles, or nymphs, carry a pair of gills on each of their first 7 abdominal segments (A1-7). These gills are flat, paddle-like organs joined to the abdomen through a thin hinge (Figure 1A). The gills on A1-6 have two lamellae each and are motile, while the gills on A7 are mono-lamellated and non-motile (21). Gills are tracheated and thought to be involved in gas exchange/respiration (e.g. (22–24); but see (25)) (Figure 1 A-C), in osmoregulation through specialized chloride cells (26, 27), and in chemosensing (28). Gill regeneration in *Cloeon* nymphs was reported more than a century ago (29). However, *Cloeon’s* regenerative capacity extends to other appendages, including legs, antennas, wing rudiments and terminal cerci (20).

**Figure 1.**
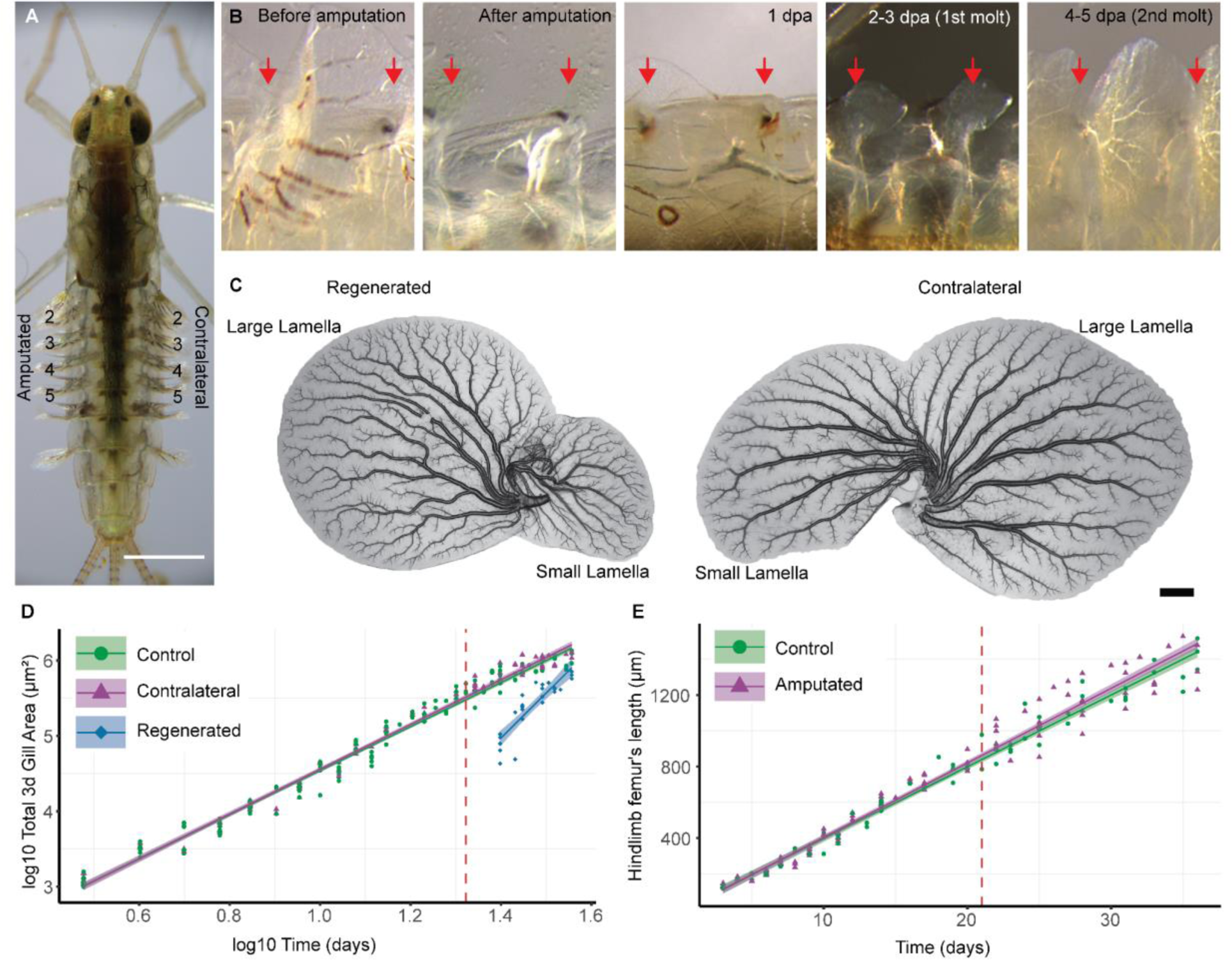
Accelerated growth of regenerating gills. **(A)** Female nymph, numbers point gills that were amputated in the experiment on the left and the contralateral ones on the right. Scale bar: 1mm (**B**) Dorsal view of third segment gill regeneration after total amputation. Anterior to the left and distal to the top. From left to right: segment before and after amputation, 1 day post amputation (1 dpa) shows a melanization clot formed in the abdomen where the gills was connected to the body, moreover the trachea that connected the gill to the tracheal system has disintegrated. Fourth panels show a regenerated gill that appears after molting once post amputation, approximately 2 dpa. After a second molt, approximately 4-5 dpa regenerating gills already have a visible tracheal tree. Red arrows indicate the point where the gills (3d and 4th) connect to the abdomen. (**C**) Whole mount gills from a last nymphal instar. Scale bar: 100 µm. Note the gill’s large and small lamella. (**D**) Linear models of control (green), contralateral (purple) and regenerated gills (blue) representing log10 Area vs log10 Time (R2: 0.9893, p-value: < 2.2e-16. Regenerated gill estimate: 1.86424 p-value: 1.55e-09). (**E**) Linear model of the hindlimb femur mean length (between left and right for each individual) over time (R2: 0.9735, p-value: < 2.2e-16. No significant differences between groups).

There are two particular aspects of gills we found of special interest. One is that gills suffer frequent autotomy (*sensu* Maginnis (6)): in our cultures we observe that gills often detach from the body and remain trapped in the shed cuticle during molting (20). This amputation happens at the base of the gill. After such amputation, a normal-looking gill regenerates within 5-9 days (Fig. 1B, C and [13]). Given the functional importance of gills for respiration, osmoregulation and chemical sensing, this regenerative capacity is likely to be important for survival. Another interesting aspect is the relation between gills and wings: the abdominal gills of mayflies have been suggested to be serially homologous to wings ((30–33) reviewed in [26]), a relationship that has been recently supported by transcriptomic analyses (28).

In this paper we shed light on these questions by describing the dynamics of gill regeneration in terms of morphology, cell and transcriptional dynamics in *Cloeon dipterum* nymphs. Our study extends the evolutionary range of insects where regeneration has been probed using molecular approaches, spanning ∼400 million years of insect evolution (34, 35). Our transcriptomic analysis identifies processes, pathways and genes already known to be involved in the regeneration of appendages in other insects, but also molecules and processes not previously associated with insect limb regeneration. We tested the functional role of orthologues of some of these genes, including components of the proteasome, the extracellular matrix and the activin pathway, in *Drosophila* larval/pupal wing regeneration. These experiments confirm the involvement of these candidates in regeneration and identify new regeneration-related mechanisms. The conservation of regeneration mechanisms between flies and mayflies supports the idea that such mechanisms may be widely conserved, at least within insects.

## RESULTS

### Accelerated growth of regenerating gills

Previous studies in the ladybug *Coccinella septempunctata* showed that these insects are able to regenerate their legs, but with a developmental cost which results in larger individuals with developmental delays (36). By contrast, the stick insect *Sipyloidea sipylus* reduces the size of its wings upon leg regeneration (37). Thus, our first experiment asked how the dynamics of regeneration compared to normal gill development, and whether gill regeneration had an impact on organismal growth. To address these questions, we separated sibling nymphs into two batches. The first was a control group raised in standard conditions (see Materials and Methods). In the second, we amputated gills on the left side of segments A2-5, at their base, at 21 days post hatching (dph). The non-amputated contralateral gills served as internal controls in the operated individuals (Figure 1 A-C). We then followed the growth of individual nymphs and collected the shed exuviae at each molt, on which we could measure the growth trajectories for each nymph until metamorphosis (note that following metamorphosis aerial adults lose the gills). We measured the areas of both the large and the small gill lamellae in control, regenerating and contralateral gills (Fig. 1D, suppl. Dataset S1). In addition, we measured the length of the femur from the third thoracic segment (“hindlimb”) as a proxy of organismal size in the control and operated groups (Fig. 1E, suppl. Dataset S1).

When we plotted the area of growing gills versus time, we observed that growth followed a logistic curve, with an early acceleration, an intermediate phase with an approximately constant growth rate and a deceleration towards the end of development (Suppl. Fig. 1). When plotting the logarithm of the area, these growth trajectories showed a linear relation with time, which allowed us to compare them easily with the growth trajectories of the regenerating and contralateral gills of operated nymphs. This comparison showed that the growth rate of regenerating gills is significantly higher than that of their contralateral controls, while the growth rate of these latter gills is indistinguishable from the rate of unoperated controls (Fig. 1D and Suppl. Fig. 2 A,B). Despite their accelerated growth, however, at the end of the nymphal period the area of regenerating gills remained slightly smaller than that of contralateral or unoperated controls (Suppl. Fig. 3A), likely because the time left for regeneration from 21 dph to metamorphosis was not sufficient for the regenerating gills to fully catch up (Suppl. Fig. 2A). When we compared the overall body growth of control and amputated individuals, using femur length as a proxy or non amputated gills from segments 1, 6 and 7, we observed that their growth trajectories overlap and reach approximately the same length at metamorphosis (Fig. 1E and Suppl. Fig. 1B; 2C,D and 3B). Our experiment showed that regenerating gills increase their growth rate without any noticeable effect on the overall growth of the injured animal.

### Faster growth is due to increased rates of cell proliferation and cell expansion

The growth of the gill area can result from cell proliferation, cell area increase, or both. In order to determine whether an increased cell proliferation rate contributed to the faster growth of regenerating gills, we repeated the experiment described above, but sacrificed the nymphs at specific time points (molts) and counted the total number of cells in each gill using DAPI staining (Fig. 2 A-A’ and Suppl Fig. 4 B; see Materials and Methods). In this experiment we corroborated the results presented earlier (Suppl Fig. 4 A) and observed that the proliferation rate is higher in regenerating gills than in controls (either contralateral gills or those from non-operated nymphs) (Fig. 2C, Suppl. Dataset S2). In order to determine whether growth resulted from localized proliferation in a “growth zone” or rather proliferation was dispersed within the regenerating gill, we pulse-labeled S-phase cells using EdU injection (see Materials and Methods). At 2-3 and 4-5 days post amputation (dpa) gills showed high and uniform density of EdU-positive cells (Fig. 2C, F, and G) consistent with a dispersed proliferation with a proportion of EdU-positive nuclei to total nuclei of 10-20%. By 6-7 dpa, the regenerated gills had reached almost contralateral size and trachea, chloride cells and margin bristles have already differentiated. These gills showed a much lower density of EdU cells, which were located mostly along the trachea of the large lamella and in the epithelium of the small lamella (Fig. 2H, Suppl. Figure 4C, and Suppl. Dataset S2), a result that is in agreement with the slower pace of gill growth towards the end of the regeneration process (Fig. 2C, Control).

**Figure 2.**
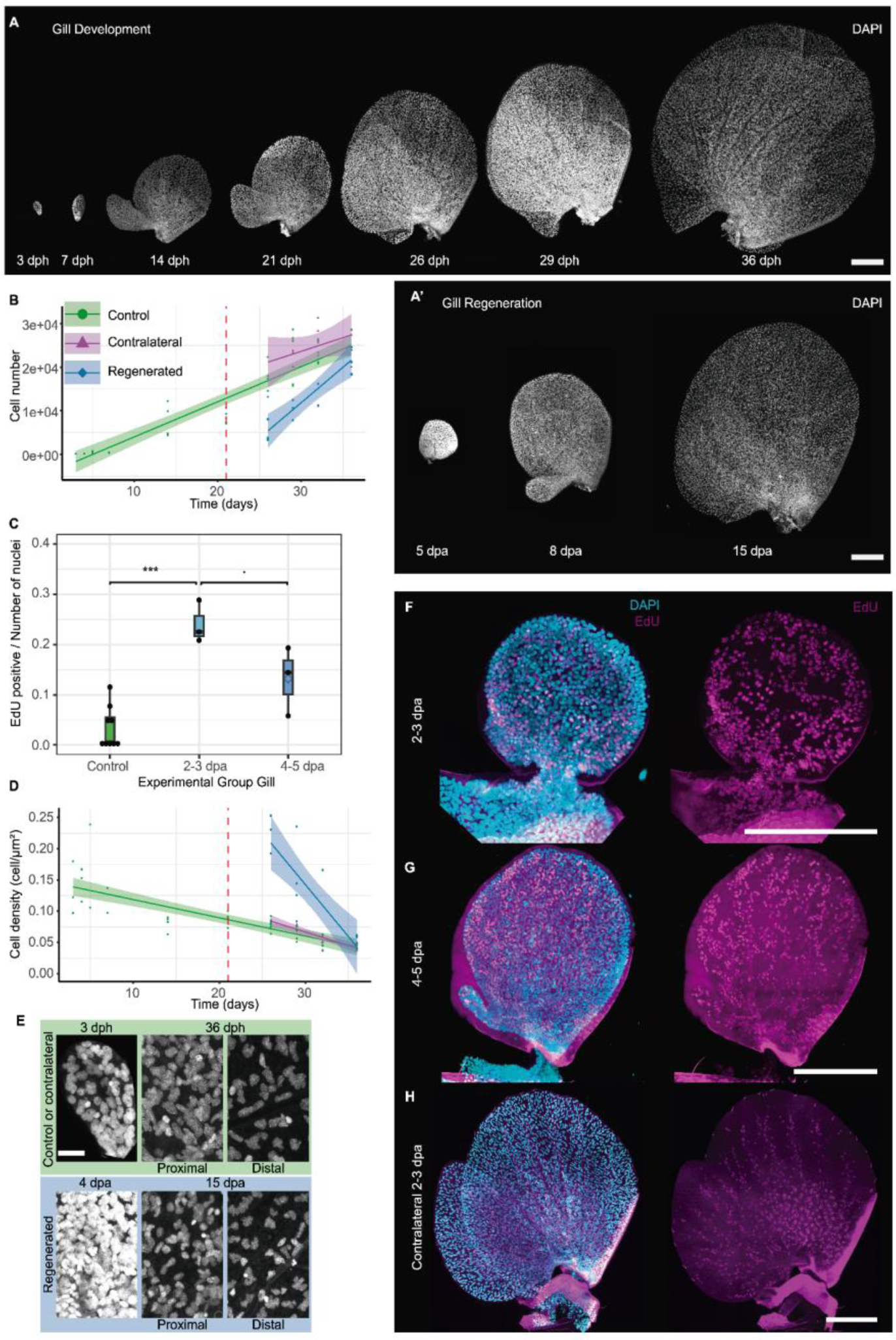
Accelerated proliferation and cell growth in regenerating gills. (**A**) Maximal z projection of DAPI (4′,6-diamidino-2-phenylindole) stained gills during development (3, 7, 14, 21, 25*, 29* and, 36* days post hatching (dph). (**A’**) regenerating gills at 4, 8, and,15 days post amputation (dpa) which corresponds to 21*, 25* and, 36* dph respectively. (**B**-**D**) Linear models of control (green), contralateral (purple) and regenerated (blue) gills representing (**B**) Number of nuclei over time (R^2^: 0.8098, p-value: < 2.2e-16. Regenerating gill estimate 826.27 p-value: 0.01517). (**C**) Boxplot representing number of EdU positive cells over the total number of nuclei for control gills in green have a mean of 3.35% EdU positive cells, regenerating gills at 2-3 dpa of 24.08%, and regenerating gills 4-5 dpa of 13.19%. Asterisk marks significant differences in the mean value using t-test comparison between groups. (**D**) Cell density (number of nuclei divided to total area) over time (R2: 0.7205, p-value: < 1.086e-15. Regenerated gill estimate 0.0137385 p-value: 1.69e-08). (**E**) Close up from panel (A) of the first gill in development (3 dph) and regeneration (4 dpa), and a fully developed gill of LNI contralateral and regenerated, from the proximal and distal part. Control panel in green and regenerating in blue. Scale bar 10μm. (**F**-**H**)Confocal images of all nuclei (DAPI staining in cyan) and the positive cells of 24h EdU incorporation assay (magenta) for gills of different regenerated stages (**F**). First molt after amputation (2-3 dpa) (**G**), second molt after amputation (4-5 dpa) (**H**), and a contralateral gil at (2-3 dpa). Scale bars 100 μm.

We also observed that during gill growth cell density decreased, suggesting that an increase in cell area also contributed to the overall growth of the gills (Fig. 2D, E). The decrease in density is observed mostly in the distal part of gills as they become flatter (Fig. 2E and Suppl. Fig. 4 B). Early regenerating gills had a higher cell density than control gills, even at the earliest stages, but by the end of regeneration the average cell density was similar in regenerated and control gills (Fig. 2D, E, and Suppl. Fig. 4 B). From these results we concluded that, during regeneration, both the rates of cell proliferation and cell expansion are higher in regenerating than in control gills.

### Transcriptome profiling of regenerating gills revealed changes associated with metabolism, cell proliferation and signaling pathways

To identify genes and pathways potentially involved in the regeneration process, we profiled the transcriptomes of regenerating gills (amputated at 21 dph) immediately after the first molt post-amputation at 2-3 dpa (sample “Reg1”, Fig. 1B) and at 4-5 dpa (sample “Reg2”, Fig. 1B). Then, we compared the transcriptional profiles of these regenerating gills with the profiles of their contralaterals (samples “Con1” and “Con2”, respectively) (see materials and Methods for details).

At 2-3 dpa, which represented the earliest time at which a gill rudiment is visible after amputation, we found 2,232 genes that were differentially expressed in regenerating gills (Reg1) relative to controls (Con1). From this set of regulated genes, 338 were upregulated and 1844 were downregulated (Dataset S3). To obtain a global view of the processes that were affected, we carried out a Gene Ontology enrichment (GO) analysis, using the functional annotations of Drosophila orthologs ((28); see Methods) (Fig. 3 A, D, G). The GO terms associated with downregulated genes (Dataset S3) were characteristic of differentiated cell types, such as transmembrane ion transport for osmoregulatory chloride cells, or synaptic organization/neuropeptide signaling for sensory cells, indicating that at this stage regenerating gill cells were less differentiated than those of the contralateral gills. The complementary set of upregulated genes (Dataset S3), even though fewer, included genes associated with mitosis and chromosomal dynamics (e.g mitotic cell cycle/sister chromatid separation [*PCNA, Mcm, RfC3* and *RfC4*, cyclins and cyclin-dependent kinases], chromosome attachment to nuclear envelope [*Nup62*], and DNA repair [*Ogg1*, *Rpr1* and *spn-A*]. Also, various histone genes coding for 4 out of five histones classes (His1, 3 genes for His2, His2A/B/H3, 3 genes for His3 and 3 genes for His4) exhibited upregulation (with none displaying downregulation), and *Nap1*, a core histone chaperone involved in histone nuclear transfer and chromatin assembly (38), was also upregulated. These transcriptional changes are indicative of increased chromatin synthesis and fast mitosis, and are in agreement with the fast proliferation of regenerating cells we observed (Fig. 2C, 2F-H). During this early stage of regeneration, we also observed a notable upregulation of histone methyltransferase and deacetylase genes, while genes related to histone acetylation were downregulated. This pattern suggested the prevalence of histone methylation marks during this early proliferative stage. The upregulation of *Su(Var)3-9, Syd4-4* and *PR-Set7* could induce the methylation of H3K27, H3K9 and H4K20, respectively (39–42)(Dataset S3). The transcriptional modulation of these epigenetic regulators is likely linked to the major gene expression changes detected.

**Fig. 3.**
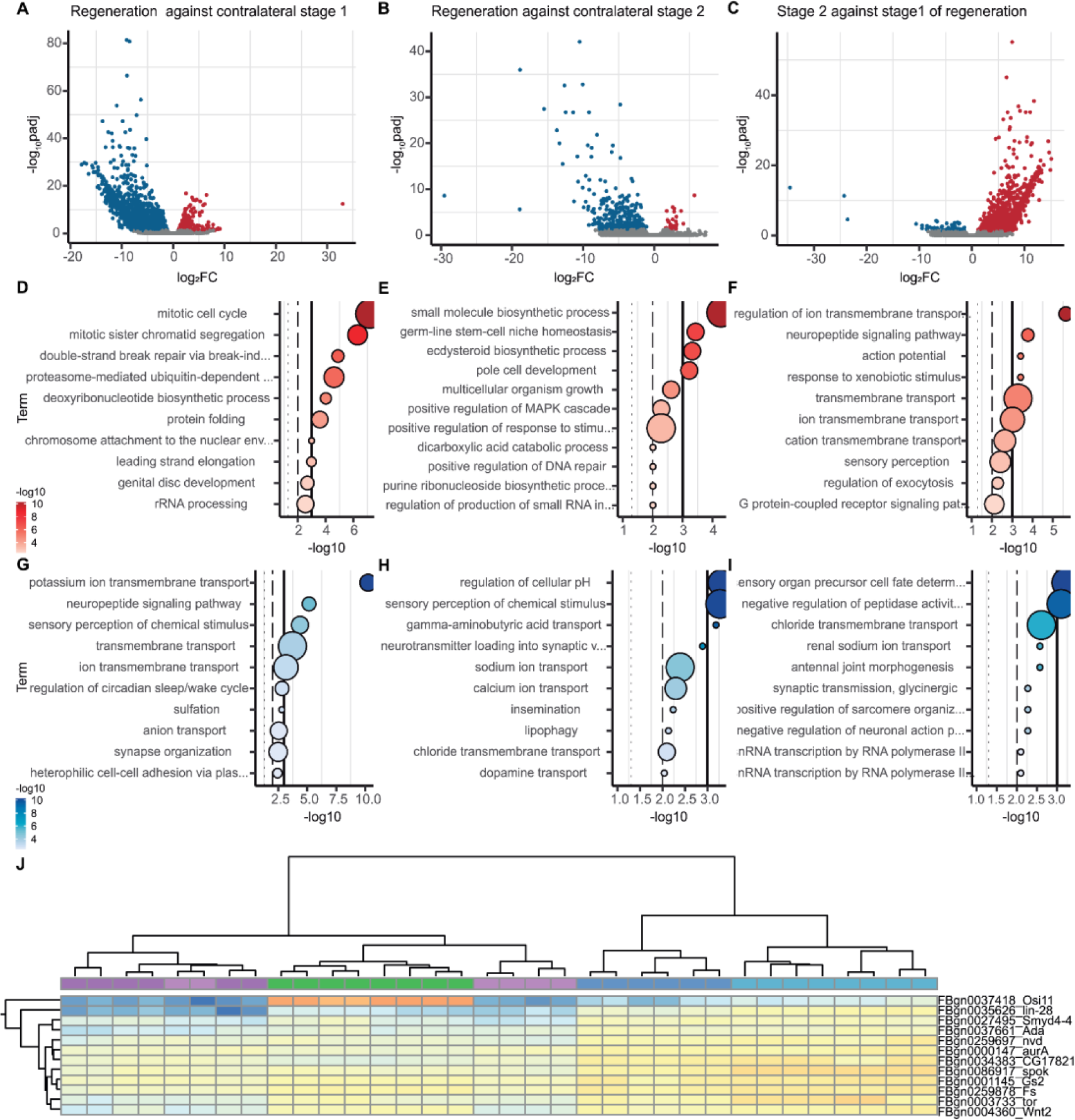
Whole gill regeneration Transcriptomics. Volcano plots for different group comparisons. Blue dots represent significantly downregulated genes, red ones significantly upregulated genes. **(A)** Stage1, regenerating gills after first molt after amputation, at 2dpa approximately, against its contralateral gills. **(B)** Stage2, regenerating gills after a second molt after amputation, at 4dpa approx, against its contralateral gills. **(C)** Regenerating gills from stage2 against regenerating gills from stage1. Dotplots representing 10 selected enriched GO terms from upregulated genes in red and for downregulated genes in blue, **(D)** and **(G)** for stage 1, **(E)** and (**H)** for stage 2 and **(F)** and **(I)** for regenerated gills from stage 2 versus stage 1. **(J)** Cluster of upregulated genes common to stage1 and stage2 that have an ortholog in *D. melanogaster* (*Osi11, lin-28, Symd4-4, Ada, nvd, aurA, CG17621, spok, Gs2, Fs, tor, Wnt2*).

In addition, two other sets of GO terms were enriched, one related to ribosomal RNA biosynthesis, presumably to sustain faster cellular growth, and the other to proteostasis (“protein folding” and “proteasome”). The latter included genes encoding components of the ubiquitination and neddylation protein-modification machinery, which were either up-regulated (e.g. *SCCRO, Cand1, TER94* y *Gint3* for “Cullin regulation and neddylation” or *APP-BP1* and *UBA3* for “E1 neddylation enzymes”) or downregulated (e.g. *Roc1a* as “part of SKP cullin ring”, and *LUBEL* and *KLHL18* for “E3s of cul3”)(Dataset S3).

When we looked for changes in components of major signaling pathways, we observed that elements of the Wnt (*wg, Wnt2, Wls, Gint3*), Hedgehog (*cubitus interruptus*), TGF-β (*Tgf-β like*) and Insulin (*Alk*) pathways are upregulated, while components of the Dpp/BMP (*dpp* and *dally*) and FGF (*breathless*) pathways are downregulated (Dataset S3). The downregulation of the FGF receptor (*breathless*) and of the transcription factor *trachealess* likely reflect the absence of trachea at this early stage (both are associated with tracheal development in *Drosophila* (43))(Dataset S3).

Globally, the transcriptome of early regenerating gills was characterized by an upregulation of genes involved in cell proliferation, changes associated with major signaling pathways, and the activation of the proteostasis machinery.

At 4-5 dpa, when differentiated structures such as margin sensillae, tracheae and chloride cells started to become visible in regenerating gills (Fig. 1B and Suppl. Fig. 5 A-L), the transcriptional profile of regenerating gills (Reg2) showed fewer differences from controls (Con2); 365 genes were differentially expressed, including 32 upregulated and 333 downregulated genes (Fig. 3 B, E, H; Dataset S4). Although regenerating gills were still proliferating faster than control gills at this stage (Fig. 2B, C, and F-H), GO terms related to cell proliferation were no longer significantly enriched. The signaling pathways that we found to be up- or downregulated at 2-3 dpa were also no longer differentially expressed relative to controls at 4-5 dpa (Dataset S4).

When the transcriptomic profiles of early and late stages of gill regeneration (Reg 1 and Reg2) were compared (Fig. 3 C, F, I), we observed a clear transition towards a more differentiated state, e.g. GO terms for neuropeptide signaling, sensory perception or ion membrane transport were enriched in the later stage (Fig 3C,F and Dataset S5). These transcriptional differences included the upregulation of several genes related to neural functions, including several Ig-containing proteins found in neuronal synapses, neuropeptides and neuropeptide receptors (Dataset S5). Ion transporters, likely linked to the differentiation of neurons and/or chloride cells, and others which in *Drosophila* are required for trachea formation and branching, were also upregulated relative to earlier stages (Dataset S5). Genes associated with these GO terms, however, were still expressed at lower levels relative to controls (Reg2 versus Con2, Dataset S4), suggesting that these regenerating gills were not yet fully differentiated. Therefore, the transition from early (2-3 dpa) to late (4-5 dpa) regenerative stages is marked by the initiation of cell differentiation and a loss of the transcriptional signature of cell proliferation.

Finally, we also identified a set of genes that were consistently up- or down-regulated in both early and late stages of regeneration (Fig. 3J and Suppl. Fig. 6). Among these, 27 genes were upregulated relative to controls (Dataset S6). These might represent a core set of genes required for the initiation and maintenance of regeneration in *Cloeon* gills. They included genes with identifiable UniProt domains and *Drosophila* homologues, as well as *Cloeon*-specific genes for which no functional annotation is available. Among those with *Drosophila* homologues, this set included genes involved in metabolism (*Gs2* (Glutamate synthase), *nvn* (Cholesterol 7-desaturase), *spok* (cytochrome P450), *CG17821* (Very-long-chain 3-oxoacyl-CoA synthase), signaling (*tor*, *Wnt2*, *Fs*), mitosis (*AurA* (aurora kinase)) and gene transcription regulation (the H3K4 methyltransferase and transcriptional repressor *Smyd4-4*)(Dataset S6).

### Genes associated with gill regeneration in *Cloeon* identify homologues required for wing disc regeneration in *Drosophila*

In order to explore whether the function of the genes we identified in our transcriptomic analysis is conserved in insects, we knocked down a set of candidate genes during *Drosophila* imaginal disc regeneration (Table 1) ((44–46) and Materials and Methods). Candidate genes were selected based on their upregulation in one or both regeneration stages. From these, we prioritized those that had been previously identified as differentially expressed during wing disc regeneration in *Drosophila* as per (Vizcaya Molina 2018). The list was also curated considering the existing literature, aiming to broaden the range of potential functions involved in regeneration. Our list included: *lin28*, encoding a small RNA-binding protein involved in stem cell maintenance in *Drosophila* (47–49), which potentiates insulin-like signaling in the intestinal stem cells by binding the InR mRNA (47); the Activin/TGFβ pathway component *Follistatin (Fs)*, encoding an Activin repressor (50, 51); the proteostasis-related *Hsp83* and *Hsp110* chaperones; *Nedd8* and *csn5*, encoding the ubiquitin like protein Nedd8 and a deneddylase which deconjugates nedd8 from cullins, respectively (52, 53); the heparin-binding extracellular ligands Miple1 and Miple2 (54); and the transcription factors *E2F1* (55) and *tfb4* (part of the RNApol_II holoenzyme (56)).

**Table 1.** List of genes functionally tested in *Drosophila*.

| Cleon<br>Gene Id | Drosophila Gen<br>name | Flybase Id | Stock | Upregulated<br>Both Stages<br>Cleon | Vertebrate<br>Orthologs |
| --- | --- | --- | --- | --- | --- |
| CD11828 | *miple1* | FBgn0027111 | Bloomington<br>id: 28067 | Yes | Midkine (MK) and<br>Pleiotrophin<br>(PTN) |
| CD11837 | *Hsp83* | FBgn0001233 | Bloomington<br>id: 32996 | First | HSP90AA1 &<br>HSP90AB1 |
| CD07674 | *Hsc70Cb_HSP110* | FBgn0026418 | Bloomington<br>id: 33742 | First | HSPH1 |
| CD06243 | *Nedd8* | FBgn0032725 | Bloomington<br>id: 33881 | No | NEDD8 |
| CD11555 | *E2f1* | FBgn0011766 | Bloomington<br>id: 36126 | First | E2f1 |
| CD02661 | *E2f2* | FBgn0024371 | Bloomington<br>id: 36674 | First | E2f2 |
| CD10126 | *nclb* | FBgn0263510 | Bloomington<br>id: 41826 | First | PWP1 |
| CD02213 | *Csn5* | FBgn0027053 | Bloomington<br>id: 62970 | No | COP9<br>Signalosome<br>Subunit 5<br>(COPS5) |
| CD09188 | *Tfb4* | FBgn0031309 | Bloomington<br>id: 65063 | First | general<br>transcription<br>factor IIH subunit<br>3 (GTF2H3) |
| CD11828 | *miple2* | FBgn0029002 | Bloomington<br>id: 55393 | Yes | Midkine (MK) and<br>Pleiotrophin<br>(PTN) |
| CD11559 | *Fs* | FBgn0259878 | VDRC Id:<br>46260 | Yes | FST |
| CD03637 | *Lin-28* | FBgn0035626 | Bloomington<br>id: 29564 | Yes | Lin28A & Lin28B |

In this assay, a transient pulse of the pro-apoptotic gene *reaper (rpr)* damages the wing primordium in a band of cells. Following this damage, the tissue regenerates during the last part of the larval period plus the pupal stage, so that adults develop wings of size and morphology similar to control flies. The function of our selected candidate genes was tested by silencing their expression during the apoptosis-induced period and checking defects in the regeneration of the wing (44)(Fig. 4A). The RNAi knock-downs against eleven of the twelve genes interfered with the regeneration in the apoptosis-induced regeneration assay (Fig. 4B), while having subtle or no effect when applied in discs without *rpr* expression (Suppl. Fig. 7A’-L’). Control wing discs, in which the pulse of apoptosis was followed by regeneration without RNAi silencing of any gene, produced morphologically normal wings in 77% of the flies, while the remaining 23% of wings showed vein fusions, or blade notching (Fig. 4B, 4C, n=90, and Suppl. Fig. 7M and N). In any case, we never observed a strong reduction of the size of the wing blade (Fig. 4C). In contrast, the defects caused when candidate genes were silenced were specific for each of them, highly penetrant and significantly stronger than controls (Fig. 4B, 4C). In particular, knocking-down *tfb4, e2f1, e2f2, miple2, csn5 and nedd8* exhibited very strong phenotypes in all wings analyzed (n=34, n=102, n=72, n=90, n=60, and n=8 respectively). Interestingly, knocking down *miple1* (n=107) did not produce as strong and consistent disruption in regeneration as *miple2*. *Hsp110*, contrary to the rest of the tested genes, had completely regenerated wings in a similar proportion to the control (72% and 77%, respectively). These results indicated that differential gene expression during *Cloeon* gill regeneration is a good predictor of genes required for wing disc regeneration in *Drosophila* and importantly, they pointed to a conserved role of some of these factors in appendage regeneration in insects.

**Figure 4.**
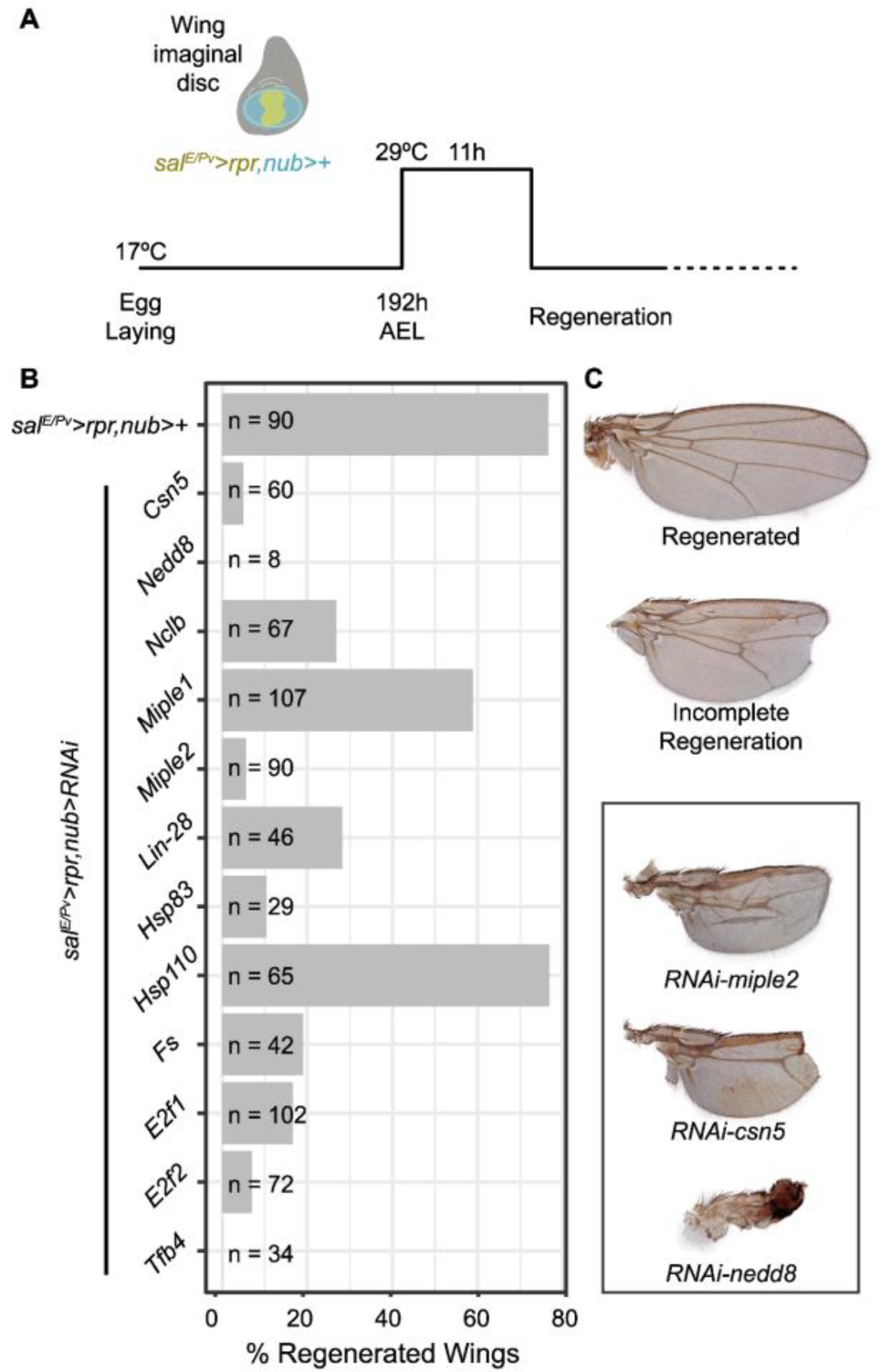
Dual transactivation experiment in Drosophila melanogaster wing disc regeneration. **(A)** Schematic representation of regeneration assays in wing disc regeneration using lexA/lexO and UAS/Gal4 systems. Reaper (rpr) expression is induced in the Spalt wing disc domain (A yellow region) after a temperature induction (29°C, which degrades for that time a temperature sensitive version of Gal80 protein (Gal80TS) which is expressed under the ubiquitous tubulin promoter. When this is performed on Drosophila larvae, 196 hours after egg laying (AEL) for 11h, cells within the spalt domain undergo apoptosis but surviving remaining cells proliferate to compensate lost cells and wings develop normally around 80% of the cases (B nub>+, C Regenerated Wing). However, when also expressing a transgene under the Upstream Activating Sequence (UAS) by using Gal4 yeast transcription factor expressed under the nubbin promoter in the wing disc pouch (A blue domain in wing disc), the wing disc may suffer defects in regeneration (C incomplete regeneration). **(B)** Histogram of percentage of fully regenerated wings with induced apoptosis and expression of RNAi for selected genes. *nub>+* indicates negative control with no RNAi induction which regenerates 80% of the cases. Number of wings examined (n) is indicated on each bar. **(C)** Microscope images of fully regenerated wings, incompletely regenerated wings, and three cases with most extreme defects in wing regeneration, induced by RNAis against *miple2*, *csn5* and *nedd8*.

## DISCUSSION

### Fast growth characterizes the early regenerative phase of *Cloeon* gills

After amputation, *Cloeon* gills regenerate through an early acceleration of growth, which is largely explained by an increased proliferation rate, accompanied by progressive thinning of the epithelium and lower nuclear density, which is indicative of an increase in cell area. The fast proliferation seen in the early regeneration stages correlates with the elevated expression of cell cycle-related genes, epigenetic modifying enzymes and components of growth-promoting signaling pathways (Fig. 3 and suppl. Dataset S3). The latter include the Insulin, Wnt and Activin pathways, and the RNA pol I regulator *nclb,* which lies downstream of the Myc and TORC cell growth pathways (57–59) (Suppl. Fig 8).

Our study does not reveal a systemic effect on the growth of the nymphs with clipped gills. Specifically, there is no developmental delay or change in the growth of uninjured gills or hindlimbs in the operated animals (Fig. 1, Suppl. Fig. 2 and Suppl. Fig. 3). However, our experiments were not designed to investigate potential systemic effects of gill regeneration and unseen effects on other organs may exist (60). Indeed, we detected a series of deregulated genes whose products could be involved in inter-organ communication due to the endocrine and paracrine functions described for their homologues in *Drosophila*. First, we detected the upregulation of *neverland* (*nvl*) and *Spok*, encoding two enzymes belonging to the Halloween gene family, known for their role in the early synthesis of Ecdysone (reviewed in (61)), while the Ecdysone receptor (*EcR*) and its co-receptor Taisman (*Tai*) were down-regulated (Dataset S3, Dataset S4). Second, we identified a group of downregulated neuropeptide and neuropeptide receptors-encoding genes (neuropeptide signaling pathway_GO:0007218, like *AstC-R2*, *CapaR*, *CCHa1*, *CCHa1-R*, and *Dh31* among others; Dataset S3-S5). These gene expression changes might be indicative of endocrine or paracrine functions of regenerating gills.

### Proteostasis and neddylation are implicated in gill/wing regeneration

During gill regeneration, many genes coding for components of the proteostasis machinery become upregulated (Fig. 3 and Dataset S3). Indeed, mounting evidence indicates that the Ubiquitin-proteasome system has an important role in stemness maintenance (reviewed in (62)), and some work has found evidence linking this system to regeneration (e.g. planaria whole body (63), sea cucumber intestine (64), axolotl limb or zebrafish axons (65)). Our work identifies an additional mechanism of proteostasis control, Neddylation, a process by which the ubiquitin-like Nedd8 peptide is covalently attached to target protein substrates (52). The major Neddylation targets are Cullins, but Neddylation affects other proteins as well, such as p53 or Ribosomal proteins (66, 67). The list of genes implicated in Neddylation which are upregulated in regenerating gills (see Dataset S3) includes *UBA3, Cand1, SCCRO*; also *Gint3 and TER94* which have been involved in Wnt signaling activation via protecting the nuclear transducer Arm from degradation by the proteasome (68). The deregulation of the Neddylation pathway during wing disc regeneration in *Drosophila* (by knocking down *nedd8* or *csn5,* an isopeptidase that de-neddylates cullins (69)) causes a dramatic impairment of wing disc regeneration (Fig. 4). What role could proteostasis be playing during regeneration? In regenerating gills cells proliferate fast, accelerating their cell cycle. This cell cycle acceleration demands fast turnover of many proteins, including cyclins and CDKs which are known substrates of the Ubiquitin/Proteasome system (70). In addition, regeneration requires cells to undergo swift state transitions, including their entry into a temporary multipotent state (71). These transitions need some degree of epigenetic reprogramming. This likely demands the regulation of chromatin-modifying enzymes, something we detected in our transcriptomic analysis (Fig. 3, Dataset S3). But beyond chromatin states, reprogramming must require the clearance of old proteins and the proper folding and stabilization of newly synthesized ones. Only through this proteasome dynamics, which would involve the action of chaperones, ubiquitination and Neddylation, the process of regeneration could take place efficiently (Fig. 4).

### Re-use of developmental programs or regeneration-specific programs?

One relevant question for understanding the mechanisms of regeneration is whether organ regeneration redeploys programs that operate during development or instead calls genes and pathways that are regeneration-specific. As mentioned above, ubiquitination and Neddylation are known to be very pleiotropic. However, the *Drosophila* wing disc regeneration assays showed that while the transient attenuation of genes such as *nedd8, csn5* or *Hsp83* do not have any noticeable effect on the development of uninjured wing primordia at larval stages (Suppl. Fig. 7), this transient attenuation dramatically impairs wing regeneration (Fig. 4). Therefore, it seems that the regenerative process is more sensitive to the alterations of this pleiotropic pathway than normal organ development. This may be the case for many of the genes and pathways we find regulated during regeneration. However, we also noted that reported *Drosophila* null mutants for *lin28, Fs, miple1* or *miple2*-which are up upregulated during *Cloeon* gill regeneration-are morphologically normal (47, 48, 54, 72), indicating that these genes play minor roles, if any, during wing development. This fact suggests that some regeneration-associated genes are indeed required specifically for wing regeneration. Our functional studies have not been comprehensive, but we believe that cases such as the ones we report here support the notion that the regenerative program of an organ utilizes genes that are not normally involved in its development. This, in turn, would imply a significant overhaul of the gene regulatory networks controlling organ development for regenerative purposes.

Finally, our functional results using *Drosophila* wing discs indicate a substantial degree of conservation for genes and processes involved in appendage regeneration of insects, since flies and mayflies diverged about 400 MYA.

## MATERIALS AND METHODS

### *Cloeon* culture and amputation procedure

*Cloeon dipterum* culture was as previously described in (20). All experiments were carried out in a room at 21 +/-1°C. Cold treatment (placement of dish on ice) was used for anesthetizing the nymphs for either gill amputation, dissection or nymph injection. Gills were amputated by pulling the gill with forceps at its base. Amputation was carried out in nymphs 21 days post hatching (dph) in which the wing pad had grown over the first half of the first abdominal segment. Gills 2-5 from the left flank were amputated except in RNAseq experiments, in which the sixth gill was also included.

### Growth dynamics: exuviae fixation, imaging, quantitation and analysis

To follow the growth dynamics of regenerating and non-operated gills, newly hatched nymphs were separated individually in 24 well plates. Wells were checked daily for shed exuviae, which were collected for fixation so that, by the last molt, all dated shed cuticles for each individual nymph had been collected. During the experiment, the water was aired by bubbling with a Pasteur pipette. Every other day ⅓ of the water was replaced. Algae were kindly provided by the Aquatic Vertebrates Platform facility at the CABD and used as food which was added daily. After about two weeks after hatching, nymphs were transferred to 50 ml beakers. Both the 24 well plates and the beakers were placed in a tray with water to make the temperature even for all individuals. Exuviae were fixed in 4% Formaldehyde for 20 min at room temperature or overnight at 4°C (without shaking). Then they were rinsed 3x in PBS and mounted in 80% glycerol in PBS. Images were taken in a Leica DM500B microscope with a Leica DFC490 digital camera. Measurements of the gill area were done manually using the polygon selection ROI tool from FIJI (73). Data analysis and plots were made in R and Rstudio.

### DAPI staining, image collection and image data analysis (TrackMate and R)

For nuclei counting nymphs were taken from different timepoints (1, 2, 4, 5, 7, 14, 21, 23, 26, 29, 32 dph and last nymphal instar (LNI)). The gut was removed and then nymphs were fixed in 4% formaldehyde in PBS overnight at 4°C with shaking. Then rinsed 4x in PBT and finally stained with DAPI (1:10.000) and phalloidin-488 overnight at 4°C with shaking. Then rinsed again 3x in PBT and 2x in PBS and then placed in 80% glycerol in PBS. The A3 gill was then mounted and imaged in a Stellaris confocal setup, using a 63x immersion objective, as z-stacks at the ALMIA platform, CABD. Nuclei in the image stacks were then counted using the TrackMate plugin (74) for FIJI. Then data was analyzed with R and RStudio.

EdU injections were done using a microinjector (Narishige IM-300) and a stereomicroscope (Leica KL300) and with the help of forceps to open the needle end to an adequate size opening (1.0 OD x0.58 ID x 100L mm, 30-0016 CAPILLARIES GC100-10 HARVARD APPARATUS). Needles need to be prepared from glass capillaries with the Flaming/Brown Micropipette Puller (or similar horizontal pullers) and conditions may change between different Micropipette Pullers and should be adjusted to the specific model.

Nymphs were injected approximately 0.2-1uL of EdU 10uM dissolved in injection solution as in (75). Nymphs were placed carefully laying on one side and all the water was removed from the Petri dish with a piece of tissue, so that the nymphs stayed put at one place for injection. Injections were performed dorsally through the space between the T3 and T4 terguites, puncturing from posterior to anterior and maintaining the needle as parallel to the dorsal cuticle of the nymph as possible. Click-it reaction for EdU was done with TermoFisher kit Click-iT™ EdU Cell Proliferation Kit for Imaging, Alexa Fluor™ 594 dye (Cat no. C10337). Most tissues of the nymphs stained well with the one click-it reaction. However, for the gills to be stained, 5 consecutive 1h-incubations in fresh reaction solution were needed. Nuclei were counterstained with either DAPI or Hoescht. Imaging was carried out in a Leica Stellaris confocal setup with 63x oil-immersion objective at ALMIA, CABD.

### RNAseq experimental design, RNA extraction, library preparation, sequencing and data analysis

For this experiment, nymphs were reared as described previously. At day 21 post-eclosion, the second to the sixth left gills were amputated, and individual nymphs were placed in separate plastic Petri dishes with algae *ad libitum*. Amputated gills at this stage were collected and this group of gills was called Con0 (Suppl. Figure 9 color green), this group was used as an external control and a proxy of a normal state of a gill within the whole inter-molting period, and this group was subsequently used in normalization later on as the within group variability was low and comparable between all groups (Suppl. Figure 9E and F). We made observations each hour for 12h during the whole regeneration process.

For stage 1 of regeneration we took nymphs that had molted once and had small regenerating gills (between 2 dpa and 3 dpa). Regenerating gills at this stage were called Reg1 (Suppl. Figure 9 color lightblue) and their contralateral gills were used as internal controls and named Con1 (Suppl. Figure 9 color light purple).

For stage 2 of regeneration we waited for the second molt, and only if in the first molt a small gill had appeared. Those nymphs that had not shown a regenerating gill after the first molt, were discarded. In this case the regeneration group was called Reg2 (Suppl. Figure 9 color dark blue) and their contralateral control gills Con2 (Suppl. Figure 9 color dark purple).

For each of the replicates we used a total of 40 individuals (around 200 gills) for each replicate. Nymphs were placed on clean water before gill amputation. Amputated gills were rinsed in a drop of cold PBS before being introduced in a prechilled Eppendorf placed on liquid nitrogen, so a fast freeze was used to preserve the RNA and help disrupt the tissue for later RNA extraction. RNA was extracted using Trizol and chloroform extraction. Biological replicates were sequenced twice and two specific samples three times, so a minimum of three biological replicates were sequenced at CNAG (https://www.cnag.eu/) using NovaSeq 6000 S1 2×50bp Paired End reads. The quality table can be found as (Dataset S8). Raw data is associated to BioProject <u>PRJNA1077402</u> with a total of 15 BioSamples and 34 SRAs objects. Quality of the read libraries were assessed using FastQC. Then reads were aligned onto the genome (GCA_902829235.1, https://www.ncbi.nlm.nih.gov/datasets/genome/GCA_902829235.1/), (GCA_902829235.1_CLODIP2_genomic.fna and GCA_902829235.1_CLODIP2_genomic.gtf files)) using STAR (76). Quality of the mapping was assessed with MultiQC. FeatureCounts was used to retrieve a count matrix of all the libraries, which had similar differences in levels of expression within groups (Suppl. Figure 9 A) and differ greatly in number of counts within and between groups (Suppl. Figure 9 B). Nevertheless, libraries cluster relatively well before normalization (Suppl. Figure 9 C and D), and after regularized normalization with DESeq2, there is really good correlation within samples (Suppl. Figure 9 F) and low variability within groups and well separation between groups in the PCA1 and PCA2 (Suppl. Figure 9 E), but for the contralateral samples (purple) that clusterized together probably due to biological similarities between those two groups. Then, DESeq2 library in R and Rstudio was used to perform differential expression analysis of the samples. Functional annotation was retrieved from available data at NCBI [21] and Uniprot (https://www.uniprot.org/). To retrieve GO annotations for *Drosophila melanogaster* ortholog genes BioMart was used. TopGO was used for gene ontology (GO) enrichment analysis with *Drosophila melanogaster* (dmelanogaster_gene_ensembl) functional annotation. Heatmaps were generated on normalized counts using “median of ratios normalization”. R libraries used in this analysis: AnnotationDbi, org.Dm.eg.db, GO.db, biomaRt, DESeq2, pheatmap, Rgraphviz, tidyr, dplyr, tibble, svglite, ggplot2 and stringr.

### *Drosophila* dual transactivation experiment

The system used is a dual Gal4/LexA transactivation system by which one of the transactivations transiently induces cell death in a stripe within the developing wing (encompassing the *sal* expression domain) by the expression of the pro-apoptotic gene *reaper (rpr).* The second transactivator system allows gene expression in a larger domain (defined by *nub* expression) that includes the death-induced stripe. Therefore, gene silencing in the cells of the wing primordium that have not been forced into apoptosis tests the effect of the gene of interest in their capacity to regenerate the dead, missing cells and the final wing. This system also evaluates the effect that the transient manipulation of the gene of interest has on wing development, thereby allowing to discriminate between the effects on development from the effects on regeneration. This system has been described previously in [29].

The specific genetic strains used were: *sal^E/Pv^-LHG* and *LexO-rpr* strains for genetic ablation. The LHG is a modified version of lexA that contains the activation domain of Gal4 separated with a hinge construct. This form is suppressible by *tubGal80^ts^* (77). The Gal-4 line used was *nubbin-Gal4* (*nub>*), which is expressed in the entire wing pouch. As both the LHG and GAL4 are suppressible by Gal80^ts^, the expression of *rpr* and the Gal4-driven UAS can be simultaneously controlled.

*w; nub-Gal4,lexO-rpr; sal^E/Pv^LHG,tubGal80^ts^*males were crossed to UAS-*RNAi (gene of interest)* virgin females. Crosses were maintained at 17°C. Synchronized 6h egg laying collections (+/-3h difference) were maintained at 17°C until day 8 after egg laying, when cultures were shifted to 29°C for 11h. This shift at 29°C inactivates the Gal80*^ts^* thus releasing both the LHG and the Gal4 transcriptional activators. After this 29°C pulse cultures were returned to 17°C until flies emerged. Emerged adults carrying both *sal>rpr* and *nub>RNAi (gene of interest)* were fixed in a mixture of Glycerol:Ethanol (1:2) for 24h. Wings were then dissected in distilled water, then rinsed in ethanol 100% and mounted in 6:5 lactic acid:Ethanol and imaged under a compound microscope. Wing patterning defects were then scored.

## ACKNOWLEDGEMENTS

We thank Alejandro Campoy for advice on image analysis and quantification and the CABD Advanced Light Microscopy Facility and Image Analysis (ALMIA) platform; Rocio Morales-Falcon, Ana Fernández-Miñan and the CABD Aquatic Vertebrates and Functional Genomic platforms; We also thank Erika Bach for the *Drosophila* UAS-Follistatin RNAi line, Michalis Averof, Jordi Solana, and David Buchwalter for comments on the manuscript, and the Casares and Almudi laboratory members for fruitful discussions.

Grants: P20_00041, Junta de Andalucía (Spain); PID2021-122671NB-I00, AEI/MICINN (Spain) to FC. PID2020-116041GB-I00, AEI/MICINN (Spain) to IA.

Genes selected for knockdown in *Drosophila* imaginal wing disc regeneration. First column has *Cloeon dipterum* gene ID, second the gene name for the ortholog in *Drosophila*, third the flybase id for the gene. Fourth column contains the ID of the fly stock with the UAS RNAi. Fifth column indicates if the gene is upregulated in both stages of Cloeon, just one (first or second) or none. Sixth indicates possible vertebrate homologous genes.

---

**Fig. S1.**
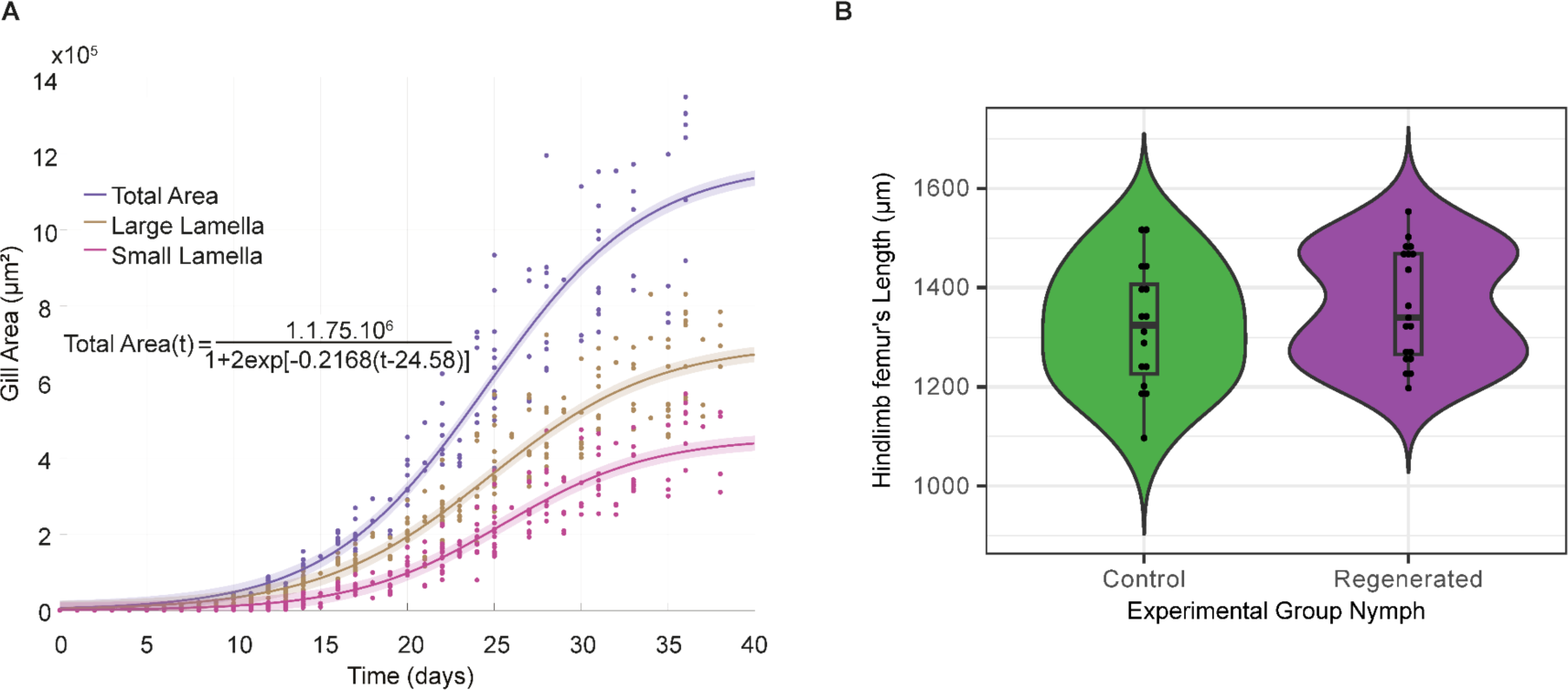
Sigmoidal fit model for gill area growth over time. Hindlimb’s femur’s length comparison. (A) Growth curve of abdominal gill 3 from ten individuals, including data and fitted curve for the area of the large (yellow) and small (magenta) lamellae and the sum of both (total area, blue). The logistic equation fitted for the total area is included. (B) Boxplot showing hindlimb femur’s length of LNI of each experimental group: control in green and operated in purple.

**Fig. S2.**
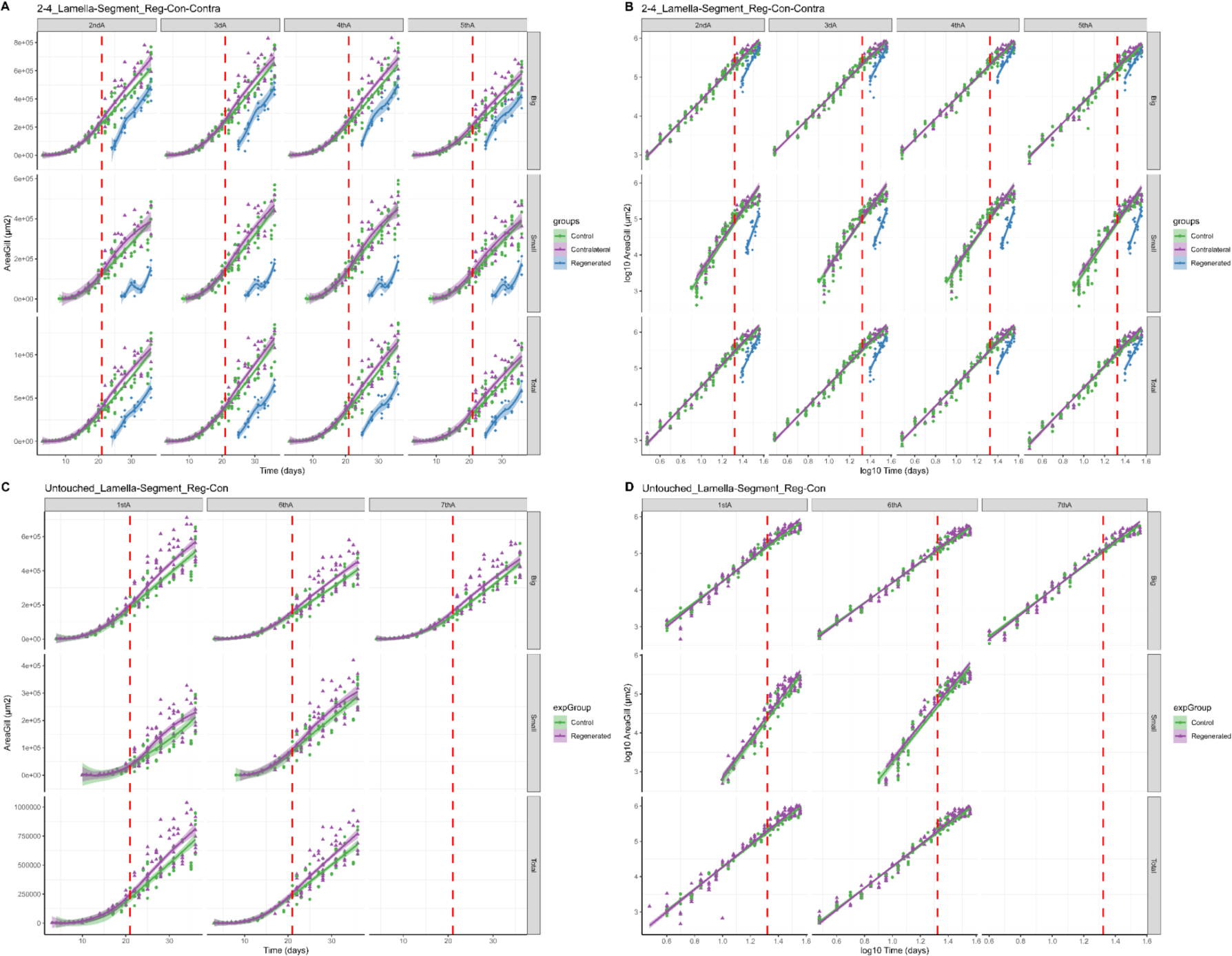
Growth curves and linear model of gills from all segments. Growth curves of Area (µm2) over time (days) for A) gills from amputated segments and gills from amputated segments. Linear models of log10 Area over log10 Time of B) gills from amputated segments and D) gills from amputated segments. Each column in each plot refers to a segment. Each row refers to big lamella (top row), small lamella (mid row) and total size (small+big, bottom row). Vertical red dashed line marks amputation day. Green represents gills from unoperated nymphs in all plots, purple represents contralateral gills for (A and B) and gills from operated nymphs from unoperated segments; and blue represents regenerating gills.

**Fig. S3.**
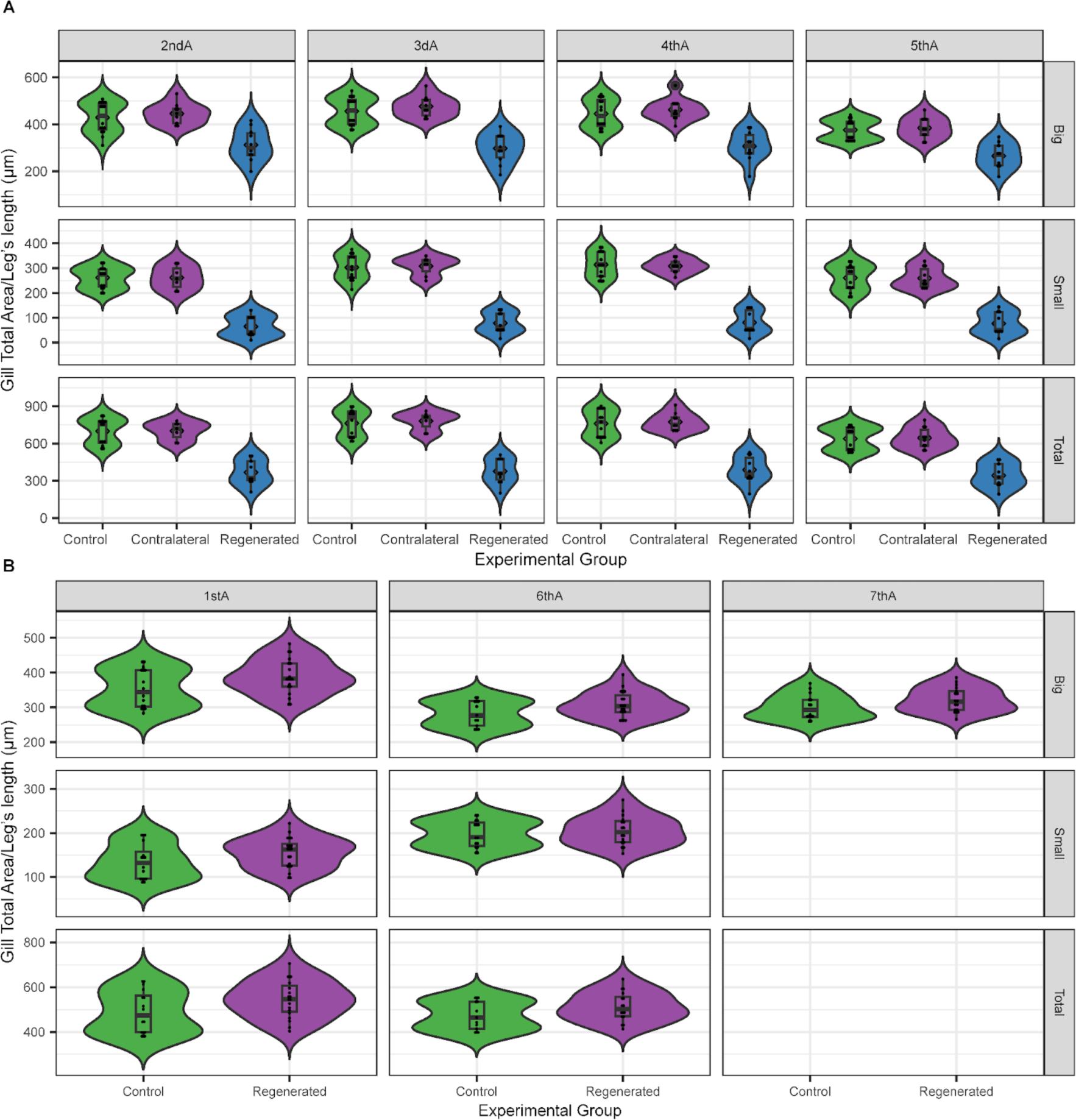
Segment gills adjusted size comparison at the last instar stage. Symmetry size within and between groups. Boxplot of adjusted gill area (Total gill area (µm22) / Mean of hindlinmb’s femur length (µm)) comparing (A) gills from amputated segments (2–5), in green those from unoperated nymphs, in purple the contralateral ones from operated individuals and in blue regenerated gills. (B) Gills from unoperated segments, from control in green and from operated individuals in purple. For each segment the areas of the large and small lamellae and their sum (total area) are plotted separately).

**Fig. S4.**
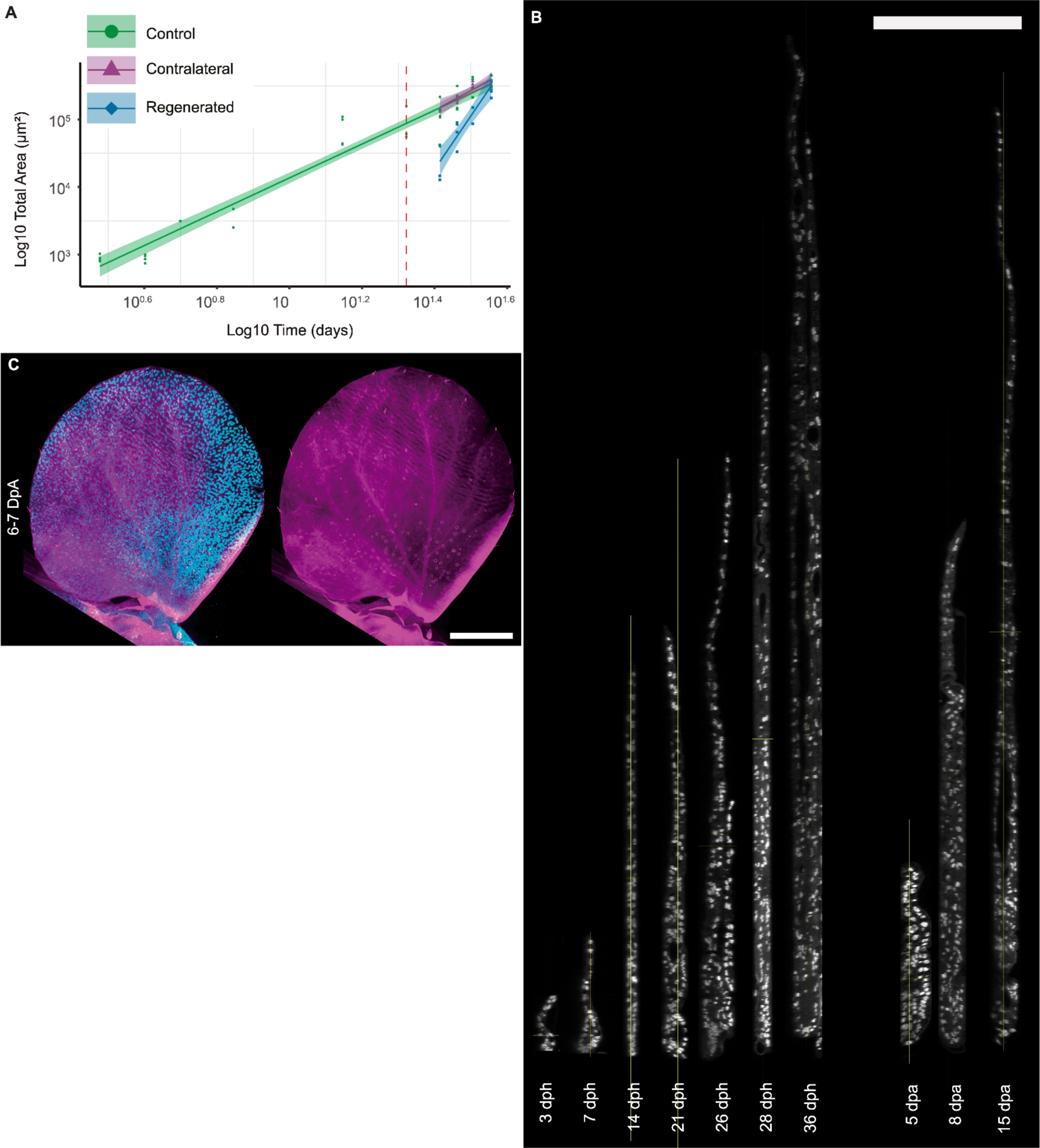
Linear model of area over time and orthogonal views of DAPI stainings and EdU 24h incorporated in regenerating gills of 6-7 dpa. (A) log10 Total Area over log10 Time (R2: 0.9414, p-value: < 2.2e-16. Regenerating gill estimate 6.43565 p-value: 1.87e-06). (B) Orthogonal views ordered by time from Figure 2 panel (A). Note that around 14 days post hatching (dph) a second lamella appears but it does not show in the cuts only for (26dph, 28dph, 36dph, 8dpa and 15dpa. In regenerating gills the small lamella appears at stage 2 (4-5 dpa). (C) Confocal images of all nuclei (DAPI staining in cyan) and the positive cells of 24h EdU incorporation assay (magenta) for gills of third molt after amputation (6-7 dpA). Scale bar represent 100µm.

**Fig. S5.**
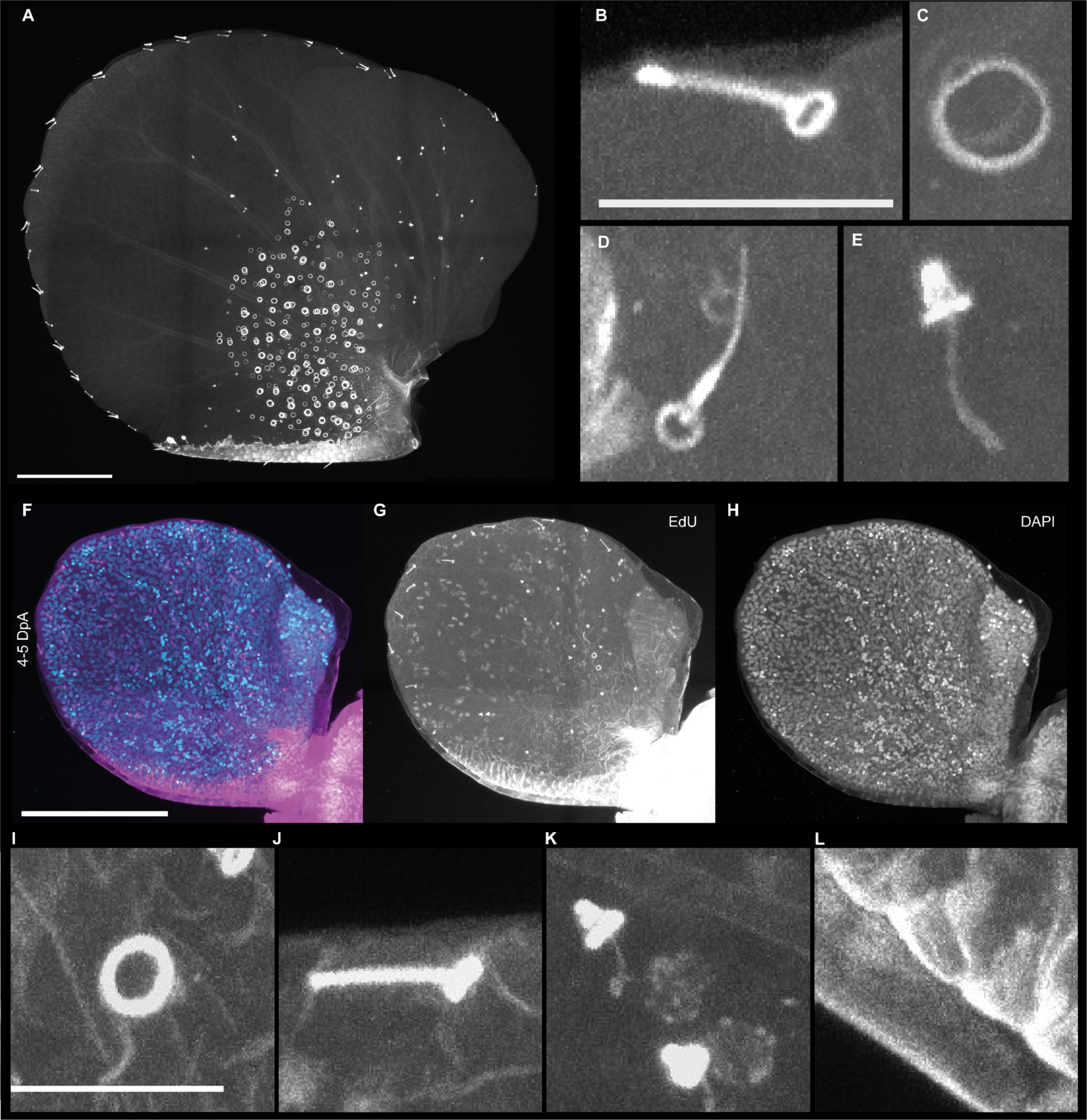
Cuticular structures of the gill as visualized using autofluorescene. A control gill of a 22 days post hatching nymph, observed under the confocal microscope using 594nm light excitation. (A) Low magnification view. “Duplicated” sensillae and chlrodide cell rims indicate the nymph was ready to shed the outer cuticle while having already synhtesized the new one, underneath. (B) margin, (D) lateral and (E) “short” sensillae. (C) cuticular rim of a chloride cell. (F-H) In some regenerating gills at stage 2 already differentiated cells can be already identified by their cuticle’s autofluorescence. Composite of EdU positive cells in magenta and nuclei stained by DAPI in cyan. (I) cuticular rim of a chloride cell. (J) margin and (K) “short”sensillae with positive nuclei and neighboring trachea. (L) Detail of the ventral rib (seen in A) underneath the old cuticle. Scale bars are 10µm for B-C and I-L, and 100µm for A and F-H.

**Fig. S6.**
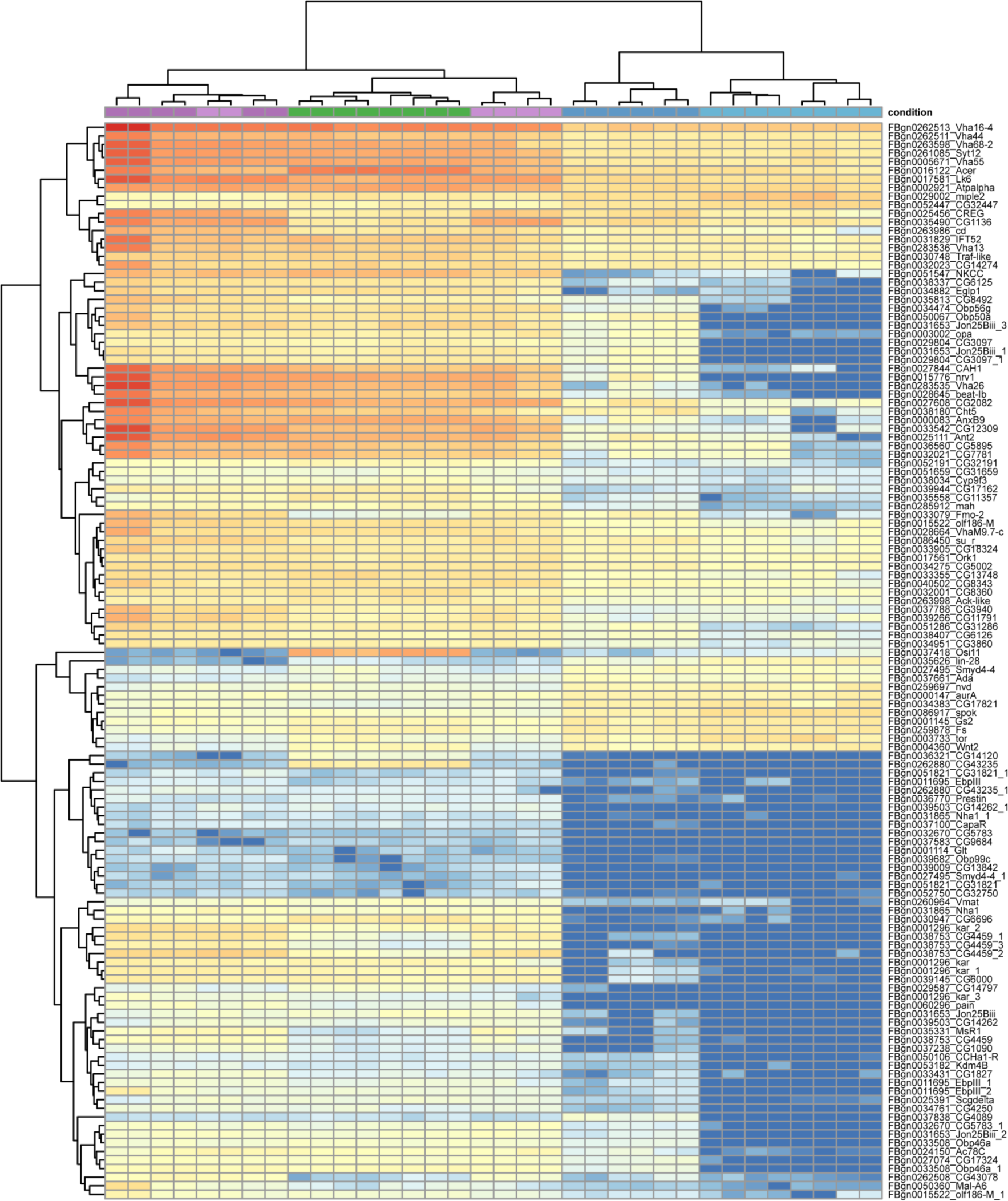
Heatmap of common DEGs in regenerating gills with orthology in Drosophila melanogaster. Heatmap of the common differentially expressed genes, with a Drosohila melanogaster possible ortholog functional annotation, between Reg1vsCon1 and Reg2vsCon2 comparisons. Each column represents a library in the RNAseq experiment. Each sample group is colored differently: regenerated gills are colored in blue (stage1 lightblue, stage2 dark blue), contralateral in purple (stage1 light purple and stage2 dark purple), and control gills in green. Each row represents the expression of a gene (counts normalized by “median of ratios normalization”). Both samples and gene expression are grouped by hierarchical clustering. All technical replicates cluster together and also all biological replicates within each sample group but for a biological replicate from contralateral stage 1.

**Fig. S7.**
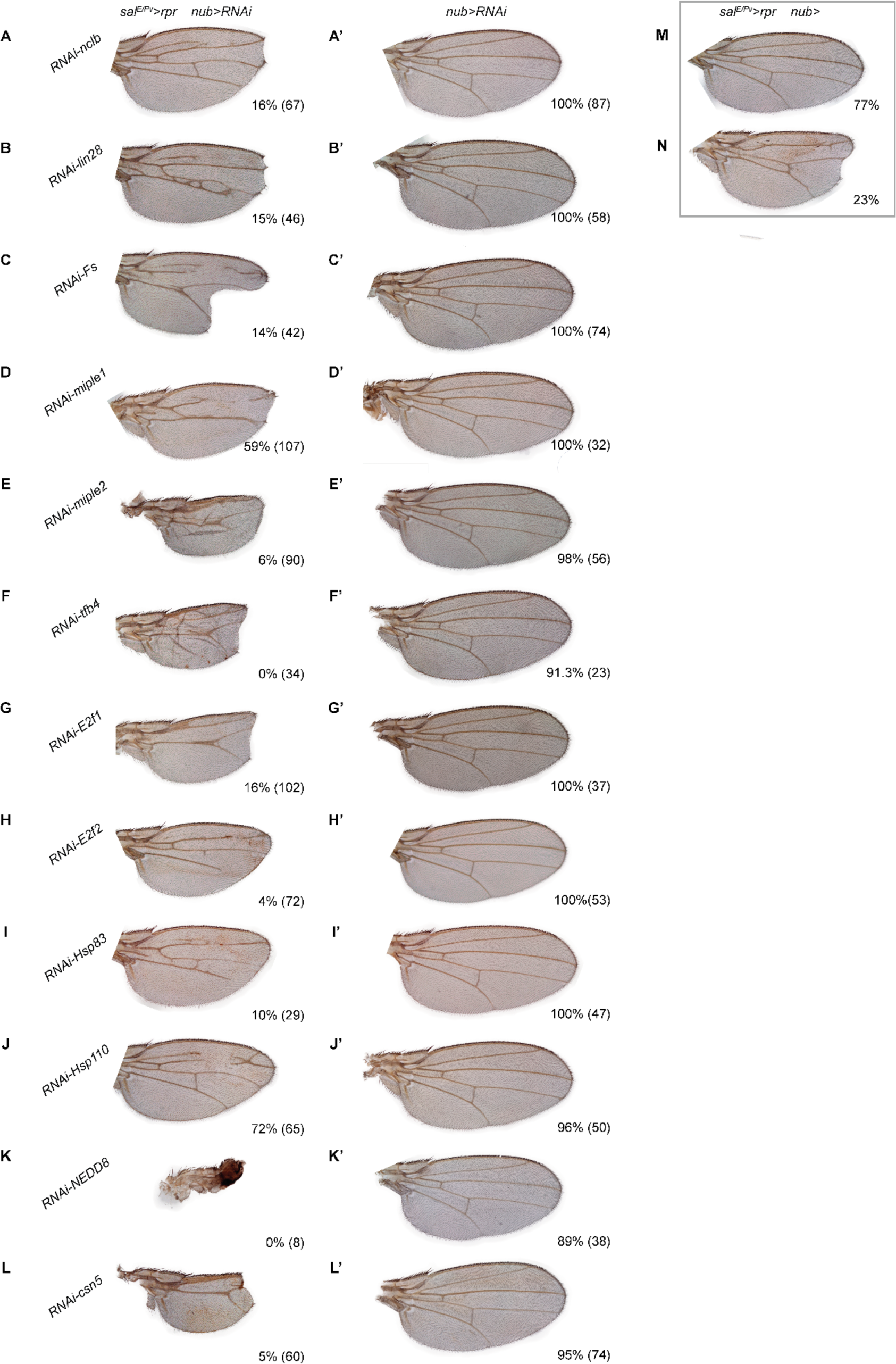
Functional screening via knockdown on Drosophila wing regeneration. Adult wings of (A-L) rpr ablated and knockdown (salE/Pv>rpr ON and nub>RNAi ON), just knockdown (nub>RNAi ON) (A’-L’). Each row of the first two columns represents a UAS-target gene RNAi: (A-A’) no child left behind (nclb) shows regenerates 16% of the time (n = 67); (B-B’) lin28 (15% n = 46); (C-C’) Follistatin (Fs) (14%, n = 42); (D-D’ and E-E’) Midkine and Pleiotrophin 1 and 2 (miple1 and miple2), (59%, n = 107; and 6%, n = 90, respectively); (F-F’) Heat shock protein 83 (Hsp83) (10% n = 29), (G-G’) Heat shock protein 110 (Hsp110) (72% n = 65), (H-H’) Transcription factor B4 (Tfb4) (0% n = 34), (I-I’ and J-J’) Elongation factor 1 and 2 (E2f1 and E2f2) (16%, n = 102; and 4%, n = 72), (K-K’) Nedd8 (0% n = 8), and (L-L’) csn5 (5% n = 60). All developmental control wings develop normally in 100% of the cases, except for (E’) miple2 (98%, n = 12), (K’) Nedd8 (89%, n = 38), and (L’) CSN5 (95%, n = 12). Control wings (rpr induction: salE/Pv>rpr ON) (M) fully regenerate 77% of the cases, with the rest of the wings showing defects as vein fusions (23%) or (N) notching in (3%). In any case, regeneration control wings do not show a significant size reduction (n = 90).

**Fig. S8.**
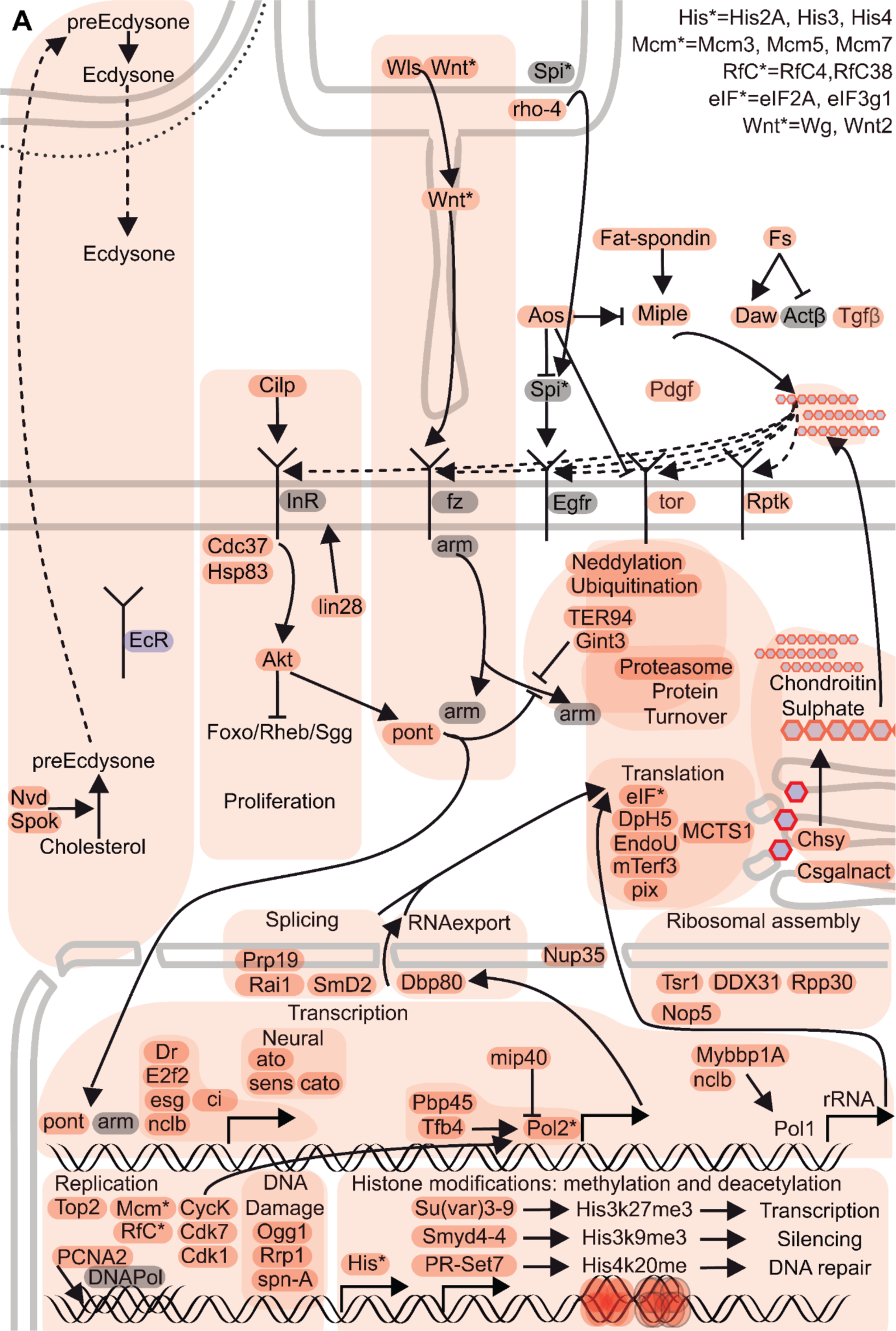
Summary of processes at stage 1 regenerating gills. Regenerating gills at stage 1. Integration of DE genes (red, upregulated; blue, downregulated; gray, not differentially expressed) along signaling pathways and processes. Insulin pathway represented by: Cloeon insulin-like peptide (Cilp), as a possible ligand; Cdc37 and Hsp83, both have been shown to bind to insulin receptor in D. melanogaster and there are two orthologs to Alk who may be repressing Foxo/Rheb/Sgg and therefore predicted to induce proliferation. Also this pathway may connect to the Wingless pathway as Alk can also activate Pontin which in turn may act protecting armadillo (arm) from proteasomal degradation and facilitating its transport to the nucleus. PreEcdysone synthesis should be favored as neverland and spookier are overexpressed. However Ecdysone Receptor is downregulated. This may result in an autonomous loss of sensitivity to Edysone while simultaneously contributing to its production. Histone expression and histone modifications (methylation and deacetylation) are activated. Other processes that are overrepresented at this stage are: DNA Replication and DNA damage; Transcription: main transcription factors, RNA polymerase II activators, RNA polymerase I activators; Spliceosome and RNA export; Ribosomal assembly, Translation, Proteasomal assembly, Ubiquitination and Neddylation regulation indicating protein turnover and proteostasis. Chondroitin sulfate synthesis. Other overrepresented signaling pathways are: Miple activated by fat-spondin and its possible binding to several receptors by binding to chondroitin sulfate. The Activin pathway is represented by Follistatin and Dawless. Argos could be repressing torso and EgfR ligands as well as activating Miple. Pointed arrows represent activation or movement and blunt-end arrows represent repression.

**Fig. S9.**
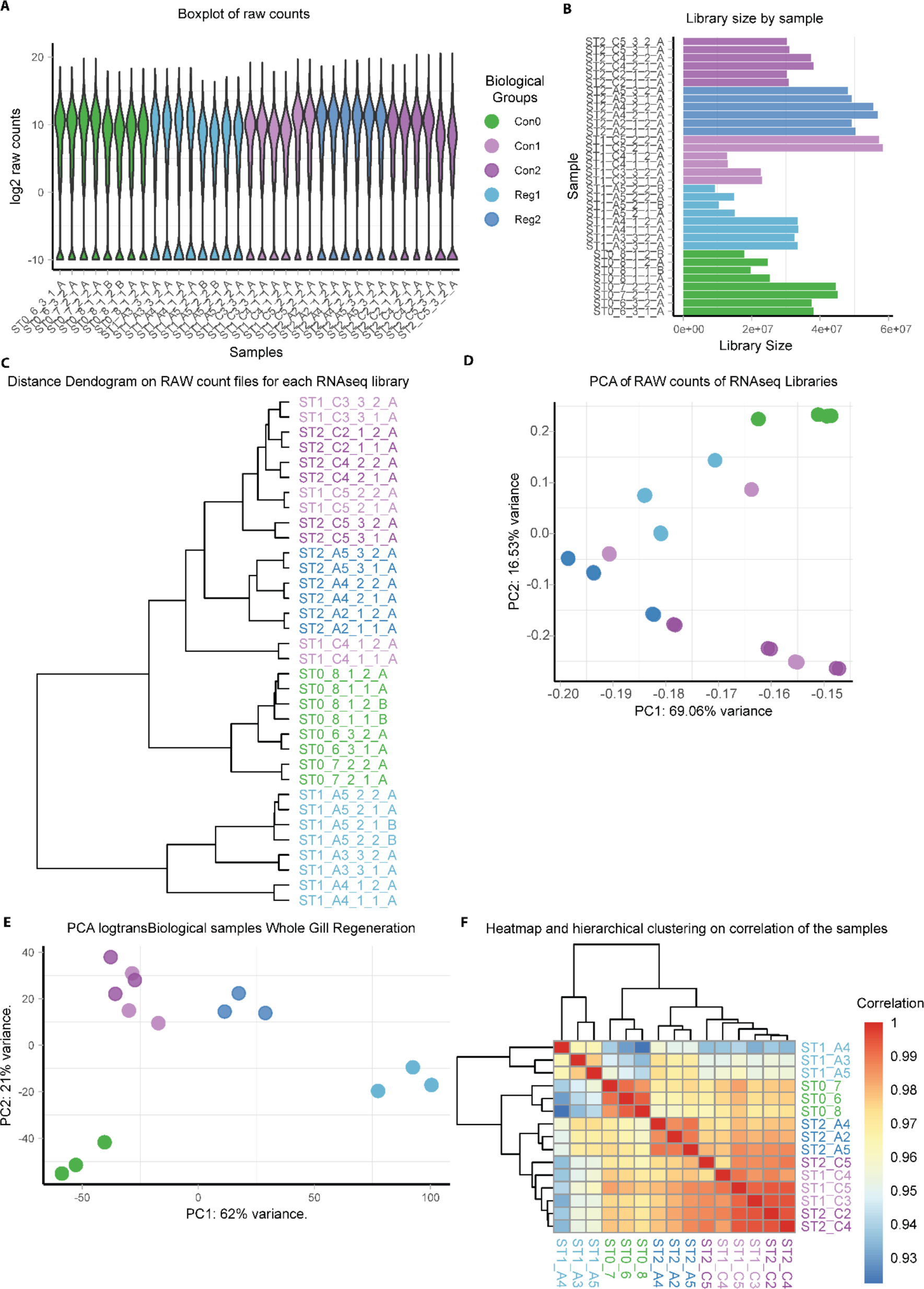
Count file quality control for each of the libraries of the RNAseq experiment before and after normalization with DESeq2. Colors of libraries or samples are according to the biological groups in the experiment (Con0 in green, Contralateral in purple and Regenerating in blue. Light and dark shades of purple and blue represent Stage 1 and Stage 2, respectively. (A) Log2 expression for all the genes in each library of the RNAseq experiment. (B) Histogram of the library size for each library in the experiment. Note that sizes differ and this will be accounted for by normalization in DESeq2. (C) Dendrogram of the distances of the raw counts for each library. All libraries clusterize by biological group but two of them belong to the same biological replicate of Contralateral stage 1. (D) Principal component analysis of the raw count file for each library. Libraries cluster by biological group but Contralateral stage 1 has a higher dispersion than the rest of the groups. (E) Principal component analysis of the count file after normalization with DESeq2 for each biological sample. Libraries cluster by biological group with low dispersion. Both contralateral stage 1 and stage 2 overlap. (F) Heatmap of the correlation between sample groups and hierarchical clustering. All samples cluster by biological group but for a Contralateral stage 2 that cluster outside a cluster of both Stage1 and Stage 2 clustering within their respective groups. Both count files before and after normalization show clusterization by biological groups.

## OTHER SUPPORTING MATERIALS FOR THIS MANUSCRIPT INCLUDE THE FOLLOWING

**Dataset S1 (separate file). Gills and femur area measurements of exuviae.**

Each row contains a sample measurement and each column (column name in “”) represents the following:

“FileName” name of the image file,

“Dates” date the sample was acquired (day-month-year),

“Appendage” which appendage the sample is (R: right hindlimb’s femur, L: left hindlimb’s femur, 1-7 are gills from segments 1-7 and the R-L indicates side and uppercase indicate large lamella whereas lowercase small lamella (for instance “1R” would be the large lamella of the right gill of the first segment)),

“Area” area measurement of length measurement in the case of the hindlimb’s femur, “symmetry” side of the sample (“Right” or “Left”),

“Segment” segment of the sample (third torathic: “3dT”, and abdominal segments from 1 to 7: “1stA”, “2ndA”, “3dA”, “4thA”, “5thA”, “6thA”, “7thA”),

“groups” (Control, Contralateral, Regenerated), “lamella” (Large or small lamella of the gill), “nymph_id” id of the nymph,

“NumDays” number of days after hatching when the exuviae was collected, “expGroup” group to which the nymph belongs to (Control, Regenerated), “mean_leg_length” mean of the left and right femur’s length of the nymph, “ID” uniq id for the measurement,

“AreaOverMLL” total area over the mean leg length.

**Dataset S2 (separate file). DAPI counts and EdU counts.**

This is a “.xlsl” file. It contains two sheets: “EdU-DAPI-Area_measurements” and “DAPI-Area_measurements”. Both sheets have quantification and measurements, each row representing a different gill. Columns represent different measurements, calculations and information of the samples.

“EdU-DAPI-Area_measurements” column description:

Name of the tiff file “FileName”, quantified number of EdU positive nuclei “EdUpositive”, automatic quantification of total number of nuclei with TrackMate “DAPI(Trackmate_unfiltered)”, number of nuclei that pass the quality and intensity threshold filtering with TrackMate “DAPI_filtered”, stage of the gill “stage” which includes (“before” = before amputation, “early” = 2-3dpa, and “mid” = 4-5dpa), gill group, “group” including (“Con”=Control, “contra”=Contralateral, and “reg”=Regenerating), time range of EdU incubation period in days post hatching (dph) “dph_24h_incubation_period”, number of cuticles the gill has (1 or two) “cuticle”, side to which the gill belongs in the nymph“side”, measured total area (large + small lamella) “area_total”, estimation of EdU positive nuclei over the total “EdU/FilteredNuclei”, estimation of cell density using nuclei over the total area “NucleiDensity(FilteredNuclei/Area)”, estimation of discarded counted nuclei with TrackMate “Nuclei/FilteredNuclei”, measured area of small lamella

“measured_area_small”, and of the large lamella “measured_area_large”, percentage of positive nuclei over the total “Percentage_EdU/FilteredNuclei”.

“DAPI-Area_measurements”

Name of the tiff file “Id”, number of abdominal segment and side the gill comes from (for instance: 3R is from the third segment right arm) “gill”, time were the nymphs were sampled in days post hatching (dph) “days”, group of nymph the gill belongs to (“CON”=Control, and “Reg”=Regenerated) “Group”, automatic quantification of total number of nuclei with TrackMate “nspots”, number of nuclei that pass the quality and intensity threshold filtering with TrackMate “filtered_spots”, days the nymph was regenerating “days_regenerating”, measured area of large lamella “area”, measured area of small lamella “area_small”, measured total area (large + small lamella) “Area_total”, group the gill belongs to (“CON”=Control, “Contra”=Contralateral, “Reg”=Regenerating) “group2”.

**Datasets S3-5 (separate files): DEseq2 and TopGO results**

All contain the same file type and order but of different differential gene expression comparisons (3 for stage 1 “Reg1vsCon1”, 4 for stage 2 “Reg2vsCon2”, and 5 for regeneration stage2 against stage1 “Reg2vsReg1”).

**“Suppl. Table 3”**

res_Reg1vsCon1.csv and Gos associated

**“Suppl. Table 4”**

res_Reg2vsCon2.csv and Gos associated

**“Suppl. Table 5”**

res_Reg1vsReg2.csv and Gos associated

They are “.xlsl” files containing 8 sheets.

The two first sheets are the result table of DESeq2 comparison, first is the complete result table and second is the filtered table by an adjusted p-value of less than 0.05. For instance for comparison Regeneration time 2 against Contralateral time 2 the sheets would be named:

“res_Reg2vsCon2” would contain all genes in the the results “res_Reg2vsCon2_padj005” would contain genes with a padj less than 0.05

This two tables contain genes in each row and each column represent the following: 1: Cloeon gene ID, 2: “baseMean”, 3: “log2FoldChange”, 4: “lfcSE”(log2FoldChange–The effect size estimate), 5: “stat” (Wald statistic), 6: pvalue, 7: padj, 8: Cloeon gene ID again “CDGENE”, possible Drosophila ortholog and flybaseID “DROME_Annotation FlybaseId”, possible Drosophila ortholog geneID “Drome_GeneId”, orthogroup ID Cloeon gene predicted protein belongs to “Orthogroup”, gene ontology ID for the possible Drosophila Cloeon ortholog “GO”, gene summary of Drosophila gene “Gene_SummaryNCBI”, Drosophila gene name “GeneName”, Drosophila gene flybaseID “FlybaseLink”, Cloeon gene predicted protein uniprot IDs (each protein from the same gene is separated by “|”) “Entry_ID_uniprot”, Cloeon gene predicted protein uniprot names (each protein from the same gene is separated by “|”)

“Protein_name_uniprot”.

The 6 remaining sheets each contain the results from the enrichment analysis of the upregulated or downregulated genes for each comparison, and for different gene ontology (GO) categories (BP: biological process, MF: molecular function, and CC: cellular component). For instance in this same comparison the sheets would be named as follows:

“Reg2vsCon2_UP_BP_allGO_topGO_analysis” upregulated Biological process “Reg2vsCon2_DOWN_BP_allGO_topGO_analysis” downregulated Biological process “Reg2vsCon2_UP_MF_allGO_topGO_analysis” upregulated Molecular function “Reg2vsCon2_DOWN_MF_allGO_topGO_analysis” downregulated Molecular function “Reg2vsCon2_UP_CC_allGO_topGO_analysis” upregulated Cellular component “Reg2vsCon2_DOWN_CC_allGO_topGO_analysis” downregulated Cellular component This tables contain GO term IDs in each row and each column represent the following: “GO.ID”; p.values for different tests (fisher and ks) performed with different methods (elim, classic and weight) in their different combinations: “p.value.elim.fisher”, “p.value.elim.ks”, “p.value.classic.fisher”, “p.value.classic.ks”, “p.value.weight01.fisher”, “p.value.weight01.ks”, annotation of the GO term “Term”, number of genes annotated with the term “Annotated”, number of genes with the term differentially expressed “Significant”, number of expected significant genes “Expected”, “Rank in Fisher.classic”; results for tests for different tests (fisher and ks) performed with different methods (elim, classic and weight) in their different combinations:“Fisher.elim”, “Fisher.classic”, “Fisher.weight1”, “ks.elim”, “ks.classic”, “ks.weight1”, term annotation and GO id separated by “_” in “Term.unique”, gene name of the significant genes appearing in the GO category “genes”.

**Dataset S6 (separate file). List of common upregulated genes in Cond1 and Cond2.**

This is a “.txt” file with each column separated by “;”. It contains 4 columns and each row represents a Cloeon gene. Column 1 has Cloeon dipterum gene ID “CDGENE”, 2 has the base mean value from DESeq2 results “baseMean”, column 3 has Drosophila possible ortholog gene id and flybase id separated by “_” “DROME_Annotation”, and column 4 contains the uniprot protein names for the possible coding predicted proteins of Cloeon genes.

**Dataset S7 (separate file). Results from Drosophila wings disc regeneration RNAi screening**

This is a table containing the genes tested in each column and each row represents a measurement. There are 6 rows:

Normal_Exp: number of normal wings in the apoptosis and knockdown experiment

Defect_Exp: number of defective wings in the apoptosis and knockdown experiment

Normal_RNAi: number of normal wings in the knockdown experiment

Defect_RNAi: number of defective wings in the knockdown experiment

TOTAL_Exp: total number of wings in the apoptosis and knockdown experiment

TOTAL_RNAi: total number of wings in the knockdown experiment

**Dataset S8 (separate files): RNAseq, quality tables.**

Xlsx file with two sheets. The first “Samples_Stat” contains the different samples sequenced at CNAG, and the second is “Libraries_Info”. Both sheets contain a description within the table.

